# RAMEN: Dissecting individual, additive and interactive gene-environment contributions to DNA methylome variability in cord blood

**DOI:** 10.1101/2025.05.08.652964

**Authors:** Erick I. Navarro-Delgado, Darina Czamara, Karlie Edwards, Maggie P. Fu, Sarah M. Merrill, Chaini Konwar, Julie L. MacIsaac, David T.S. Lin, Piush Mandhane, Elinor Simons, Padmaja Subbarao, Theo J. Moraes, Jari Lahti, Gregory E. Miller, Elisabeth B. Binder, Katri Räikkönen, Stuart E. Turvey, Keegan Korthauer, Michael S. Kobor

**Affiliations:** Edwin S. H. Leong Centre for Healthy Aging, Faculty of Medicine, University of British Columbia, Vancouver, BC, Canada; British Columbia Children’s Hospital Research Institute, Vancouver, BC, Canada; Centre for Molecular Medicine and Therapeutics, University of British Columbia, Vancouver, BC, Canada; Max-Planck-Institute of Psychiatry, Department of Genes and Environment, Munich, 80804, Germany; The Warren Alpert Medical School of Brown University; University of Alberta, Edmonton, AB, Canada; UCSI university, Kuala Lumpur, Malaysia; Department of Pediatrics & Child Health, University of Manitoba, Winnipeg, MB, Canada; Translational Medicine Program, The Hospital for Sick Children, Toronto, ON, Canada; Department of Psychology and Logopedics, Faculty of Medicine, University of Helsinki, Helsinki, Finland; Folkhäsan Research Centre, Helsinki, Finland; Department of Psychology and Institute for Policy Research, Northwestern University, Evanston, Illinois; Department of Psychology, Faculty of Medicine, University of Helsinki, Helsinki, 00014, Finland; Department of Obstetrics and Gynecology, Helsinki University Hospital, 00290 HUS; Department of Pediatrics, Faculty of Medicine, University of British Columbia, Vancouver, Canada; Department of Statistics, University of British Columbia, Vancouver, BC, Canada; Department of Medical Genetics, Faculty of Medicine, University of British Columbia, Vancouver, BC, Canada

**Author notes:** Co-corresponding authors; position determined by inverse alphabetical order,. First author (Erick I. Navarro-Delgado).

## Abstract

DNA methylation (DNAme) is the most commonly studied epigenetic mark in human populations. DNAme has gained attention in the Developmental Origins of Health and Disease field due to its gene expression regulation and potential long-term stability. Genetic variation and environmental exposures are amongst the main factors influencing inter-individual DNAme variability. However, the proportion and genomic distribution of their individual, additive and interactive effects on the DNA methylome remains unclear. Here, we introduce RAMEN, a Findable, Accessible, Interoperable, and Reusable (FAIR) framework tailored for DNAme microarrays. Using machine learning and statistical techniques, RAMEN models and dissects gene-environment contributions to genome-wide Variably Methylated Regions (VMRs), while controlling for spurious associations. To comprehensively test the power of RAMEN, we analyzed and characterized VMRs from cord blood samples from two independent cohorts (CHILD and PREDO; overall n=1,662). We identified genetics as a consistent key contributor to DNAme variability, usually in additive and interactive combinations with the environment, with genetic terms explaining the largest proportion of DNAme variance, compared to environmental and interaction terms. Operationalizing RAMEN as an R package to conduct scalable genome-exposome contribution analyses, our results highlighted the importance of genetic variation in sculpting DNAme patterns in early life.

## Introduction

The prenatal period is a time of heightened sensitivity that marks the beginning of an individual’s health trajectory. Developmental programming, a concept often studied under the Developmental Origin of Health and Disease (DOHaD) framework, describes how exposures and experiences during the prenatal period or shortly after can lead to long-lasting biological effects, potentially impacting future health outcomes^1^. At the molecular, several processes may facilitate developmental programming, including DNA methylation (DNAme). DNAme is an epigenetic mark defined by the addition of a methyl group to the DNA primarily in a cytosine-guanine (CpG) context^2,3^. Briefly, DNAme is a plausible candidate for biological embedding of environments due to its role in regulating gene activity^4,5^ and its longitudinally-stable yet environmentally-malleable nature^6–9^.

Furthermore, DNAme changes have been associated with a wide variety of environmental exposures^3,4,9^ and health outcomes^3–5^. However, the factors that contribute to shaping DNAme variation across the human genome are just beginning to be understood. Characterizing these factors is crucial to better understand disease-associated DNAme patterns, which might provide innovative avenues to refine precision medicine approaches.

Inter-individual DNAme differences in human populations are primarily, but not exclusively, associated with two factors: genetic variants^10^ and environmental exposures^11^. Single Nucleotide Polymorphisms (SNPs) associated with DNAme levels, termed methylation quantitative trait loci (mQTLs), tend to be mostly stable across the lifespan^12^ and associated with a significant proportion of CpGs in the human methylome (in the range of 1 to 48%) predominantly when they are in *cis* (reviewed in Villicaña & Bell^10^). Besides genetics, multiple associations between environmental exposures and DNAme changes have been reported. Specifically in early life, DNAme signatures have been associated with biological, nutritional, socioeconomical and pollutant exposures^11,13,14^.

In addition to the individual associations of genetics and the environment, environmental exposures could have differential influence on DNAme depending on the underlying genetic makeup. Recent studies in blood have reported gene-environment interactions (GxE) associated with DNAme levels in psychological and environmental exposures^15–19^. These interactions add a layer of complexity in the study of DNAme, since genome-wide investigation requires a larger sample size and more computationally expensive statistical methods^20^. However, characterizing these interactions offers a highly promising avenue to understand the etiology of complex human traits^21^. Overall, there is robust evidence pointing at both genetic variation and environmental exposures as sources of DNAme variation. However, integrative studies are required to dissect and estimate the individual, joint and interaction contribution of these two factors across variable regions of the DNA methylome.

Pioneering integrative studies in birth tissues have estimated that DNAme levels of Variably Methylated Regions (VMRs; genomic regions with high levels of inter-individual DNAme variability) are better explained by additive or interactive mathematical models that include genetic and prenatal environmental variables as opposed to only one of these factors^22–24^. While these studies laid the groundwork for modelling the methylome-wide contributors to DNAme variability, recent advances in study design and analysis methods now offer the opportunity to refine this framework. Briefly, previous studies measured a limited number of prenatal environmental exposures ranging from 8 to 19, which could result in an underestimation of the environment’s contribution to DNAme variability. Furthermore, studies examining the genome-exposome contribution to methylome variability could potentially improve estimations through analytical updates. Specifically, taking into account the DNAme microarray design to best capture methylome variability patterns, mitigating the effect of the imbalance in the number of variables commonly available in genome and exposome data sets to compare them on more equal methodological ground, and controlling for the spurious models expected by chance.

In this study, our main objective was to characterize DNAme variability patterns in cord blood and identify whether they were best explained by differences in genetics (G), environmental exposures (E), their independent additive effects (G+E), or their interaction (GxE) through a reproducible bioinformatic framework that addresses the limitations of previous work. To do so, we integrated genome, exposome and methylome data from 699 cord blood samples of the Canadian Healthy Infant Longitudinal Development (CHILD) study^25–27^. This cohort possesses a comprehensive characterization of prenatal exposures, capturing 94 curated prenatal environmental variables including maternal psychosocial factors, built environment, maternal nutrition and maternal health measurements. To allow for a reproducible and structured analysis of such data, we developed the R package RAMEN (*Regional Association of Methylome variability with the Exposome and geNome*) a Findable, Accessible, Interoperable, and Reusable (FAIR)^28^ tool to conduct a genome-exposome contribution to DNA methylome variability analysis. Specifically, in this paper we aimed to: 1) identify cord blood VMRs; 2) identify the model (G, E, G+E, GxE or null) that best explained the methylome variability per VMR; 3) characterize the regions explained by each model in CHILD; and 4) apply our methodology and compare results with those previously published on the Pre-eclampsia and Intrauterine Growth Restrictions (PREDO) cohort^23^.

Our results showed that genetics was a consistent main contributor to DNAme variability in cord blood, frequently in additive and interactive combinations with the prenatal environment. Using a FAIR methodology, our comprehensive characterization on how and where in the genome genetics and environmental exposures associate with DNAme variability shed light on the complex interplay between these factors during a highly sensitive developmental period.

## Results

### RAMEN provided a reproducible framework to estimate the gene-environment contribution to DNA methylation across genomic regions

We developed *RAMEN (Regional Association of DNA Methylome with the Exposome and geNome)*, an R package that provides a FAIR and user-friendly tool to dissect the factors associated with genome-wide DNA methylation variability patterns. *RAMEN* integrates matched microarray DNA methylome data, genomic variants, and exposome data, and analyzes them in a framework that can be conceptually divided into three main steps.

The first step identifies the top 10% most variable DNAme probes in a given data set based on their variance. In the second step, these highly variable probes are grouped into Variably Methylated Regions (VMRs). Modelling DNAme variability through regions rather than individual CpGs provides several methodological advantages in association studies, since CpGs display a significant correlation for co-methylation when they are close (≤1 kilobase)^29,30^. Some of these advantages include increasing statistical power by testing redundant probes only once, reducing false-positives driven by one problematic probe in a region, and improving comparability between studies that analyze the same genomic region but measure distinct CpGs due to microarray design differences^31^. In addition to canonical VMRs, defined as two or more proximal probes with correlated DNAme levels, we implemented an additional VMR category to account for the sparse and non-uniformly distributed coverage of CpGs in microarrays. Non-canonical VMRs aimed to retain genomic regions with high DNAme variability measured by single probes. This is particularly relevant in the Illumina EPIC v1 array, where most covered regulatory regions (up to 93%) are represented by just one probe. Notably, based on empirical comparisons with whole-genome bisulfite sequencing data, these single probes are mostly representative of local regional DNAme levels due to their positioning (98.5-99.5%)^32^. By taking into account the probes positioning design from the microarrays during the VMR identification step, we aimed to improve the capture of methylome variability patterns. In the last step, the individual and joint contribution of genetic and environmental variables to each VMR is estimated through machine learning and statistical techniques (**Figure 1A**; see Methods for further details).

**Figure 1.**
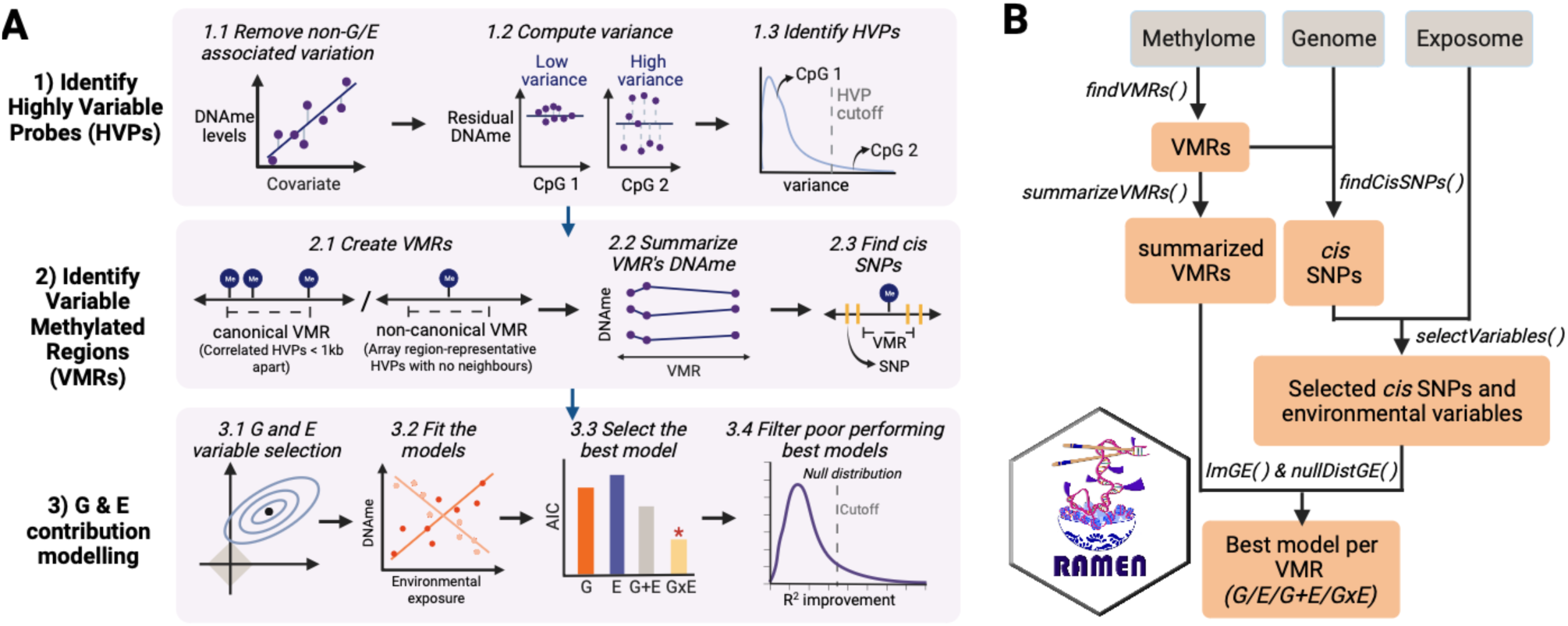
RAMEN provides a FAIR framework to model gene-environment contributions to DNA methylome variability. A) Conceptually, the first step involves identifying the 10% most variable probes in the DNA methylation microarray based on their variance (steps 1.1-1.3). Next, the HVPs are grouped into VMRs based on genomic proximity and DNAme correlation (step 2.1). The DNAme levels of each region are summarized per individual by taking the median DNAme level of the constituting probes (step 2.2). After identifying the SNPs in *cis* for each VMR (step 2.3), a machine learning algorithm is used to screen *cis* SNPs and environmental exposures and select potentially relevant variables per VMR (step 3.1). Selected variables are then used to fit G, E, G+E and GxE models (step 3.2), and the best model is chosen by comparing their Akaike Information Content metrics (step 3.3). Finally, a permutation analysis is conducted to simulate a null distribution that is used to filter out winning models that do not perform better than expected by chance (step 3.4). B) The RAMEN package implements the methodology in six functions: *findVMRs* (steps 1-2.1), *summarizeVMRs* (step 2.2), *findCisSNPs* (step 2.3), *selectVariables* (step 3.1), *lmGE* (steps 3.2-3.3), and *nullDistGE* (step 3.4). See Methods for further details.

We implemented the RAMEN framework in 6 core functions compatible with parallel computing that identify and summarize VMRs in DNAme microarrays, select G and E variables potentially associated with DNAme, identify the best model per VMR (G, E, G+E or GxE), decompose DNAme variance, and simulate a null distribution to identify non-conclusive (NC) models within each data set (**Figure 1B**).

Cord blood VMRs were enriched in quiescent chromatin, open sea, intergenic and gene body regions.

Putting RAMEN into practice, we took advantage of the CHILD cohort, which has DNAm profiles from 699 cord blood samples along with genotyping and extensive prenatal exposome characterization. After removing variation associated with concomitant variables (sex, gestational age, estimated cell type proportion and population stratification), we identified 78,569 Highly Variable Probes (HVPs; top 10% probes with the highest variance). These HVPs captured 70% of the 3,885 previously reported tissue- and ethnicity-independent human highly variable CpGs (hvCpGs) in the EPIC array^33^, as expected based in the studies’ definition of hvCpGs (top 5% variable CpGs in ≥ 65% data sets^33^). Next, we grouped the identified HVPs into regions based on probe proximity and correlation, resulting in 28,480 Variable Methylated Regions (VMRs) that were distributed roughly evenly across the autosomes (**Figure 2A**; additional details in Methods). VMRs were predominantly located in open sea, gene body and intergenic segments, and quiescent chromatin states, and, compared to the array background probe distribution, they were significantly enriched in all those categories except for gene body (FDR < 0.05; **Figure 2B**). VMRs were also primarily composed of the non-canonical type (**Figure 2C**).

**Figure 2.**
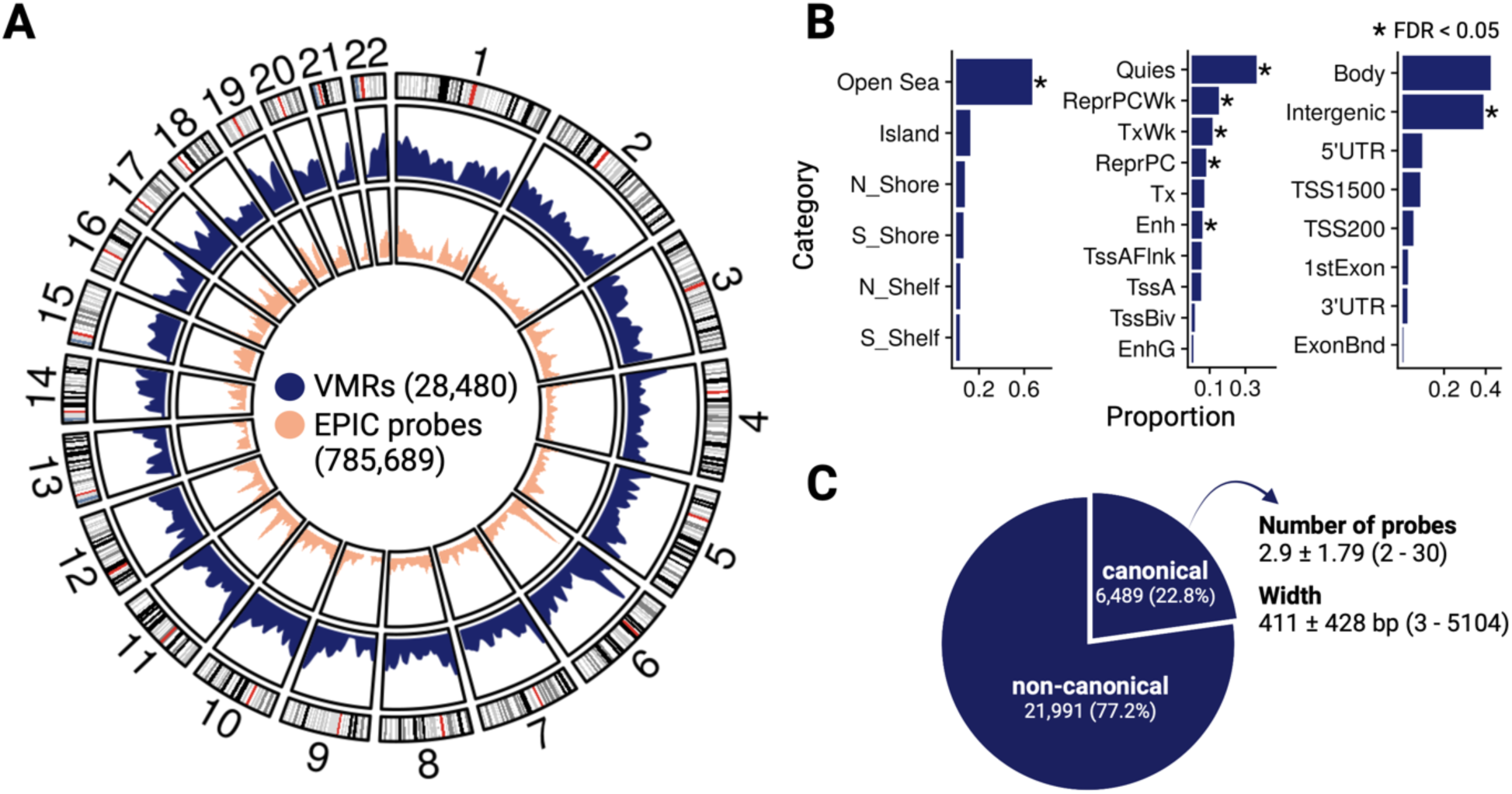
Cord blood VMR were mainly located in open sea, quiescent chromatin, gene body and intergenic regions across the genome. **A)** Density of VMRs across the genome. The background distribution of EPIC array probes is displayed in the most inner circle; circos plot was created with *circlize*^80^. **B)** Characterization of VMRs regarding CpG island context, chromatin state and gene annotation. The FDR corresponds to a one side Fisher’s test to identify enrichments in the VMR probes relative to probes in the Illumina EPIC array. For the chromatin state, only the top 10 most frequent states are showed. Definition of categories: Shore = 0.2 kb from island, Shelf = 2-4 kb from island, N = upstream(5’) of CpG island, S= downstream (3’) of CpG island; Quies - quiescent/low; ReprPCWk = weak repressed PolyComb; TxWk = weak transcription; ReprPC = repressed PolyComb; Tx = strong transcription; Enh = enhancers; TssAFlnk = flanking active transcriptional start site (TSS); TssA = active TSS; TssBiv = bivalent/poised TSS; EnhG = genic enhancers; TSS200 = 0–200 bases upstream of the TSS; TSS1500 = 200–1500 bases upstream of the TSS; 5’UTR = Within the 5’ untranslated region, between the TSS and the ATG start site; Body = Between the ATG and stop codon, irrespective of the presence of introns, exons, TSS, or promoters; 3’UTR = between the stop codon and poly A signal. **C)** Composition of VMRs regarding type, average number of probes and width.

### RAMEN reduced and balanced the number of genome and exposome variables through feature selection

Following the identification of cord blood VMRs, we modeled the contribution of the genome and exposome to their DNAme levels in four different scenarios: DNAme levels being best explained by a genetic difference (G), an environmental exposure (E), or the combination of genetics and the environment in an additive (G+E) or an interactive (GxE) manner. To model the exposome contribution, we used 94 prenatal exposome variables characterized in the CHILD cohort that encompass four main fetal exposure dimensions: maternal health (48.9%), maternal nutrition (30.9%), maternal psychosocial state (10.6%), and built environment (9.6%; **Supplementary Data 1**). To model the genetic contribution, we utilized imputed DNA microarray genotyping data and included the SNPs in *cis* of each VMR (*cis:* < 1Mb up or downstream; mean *cis* SNPs per VMR = 980.3, *SD* = 415.3, range = [0-6172]).

Using RAMEN, we observed a substantially higher number of G variables compared to E across VMRs, which could potentially bias the models towards including G factors, as the proportion of VMRs with a G association could be higher compared to E simply by chance. In addition, the high number of variables could lead to a high computational burden associated with fitting all combinations of G, E, G+E and GxE models, which were on the order of 10^9^. To balance the number of genome and exposome variables and remove uninformative variables, we implemented in RAMEN a scalable machine learning feature selection strategy based on *Least Absolute Shrinkage and Selection Operator* (LASSO) before fitting the models. After feature selection, we observed a reduction in the number of *cis* SNPs and environmental exposures considered for modelling per VMR from 980 to 16 (G) and 94 to 3 (E) on average, and a reduced difference between average G and E variables per VMR (initial: 886, post-selection: 13; **Figure 3A**).

**Figure 3.**
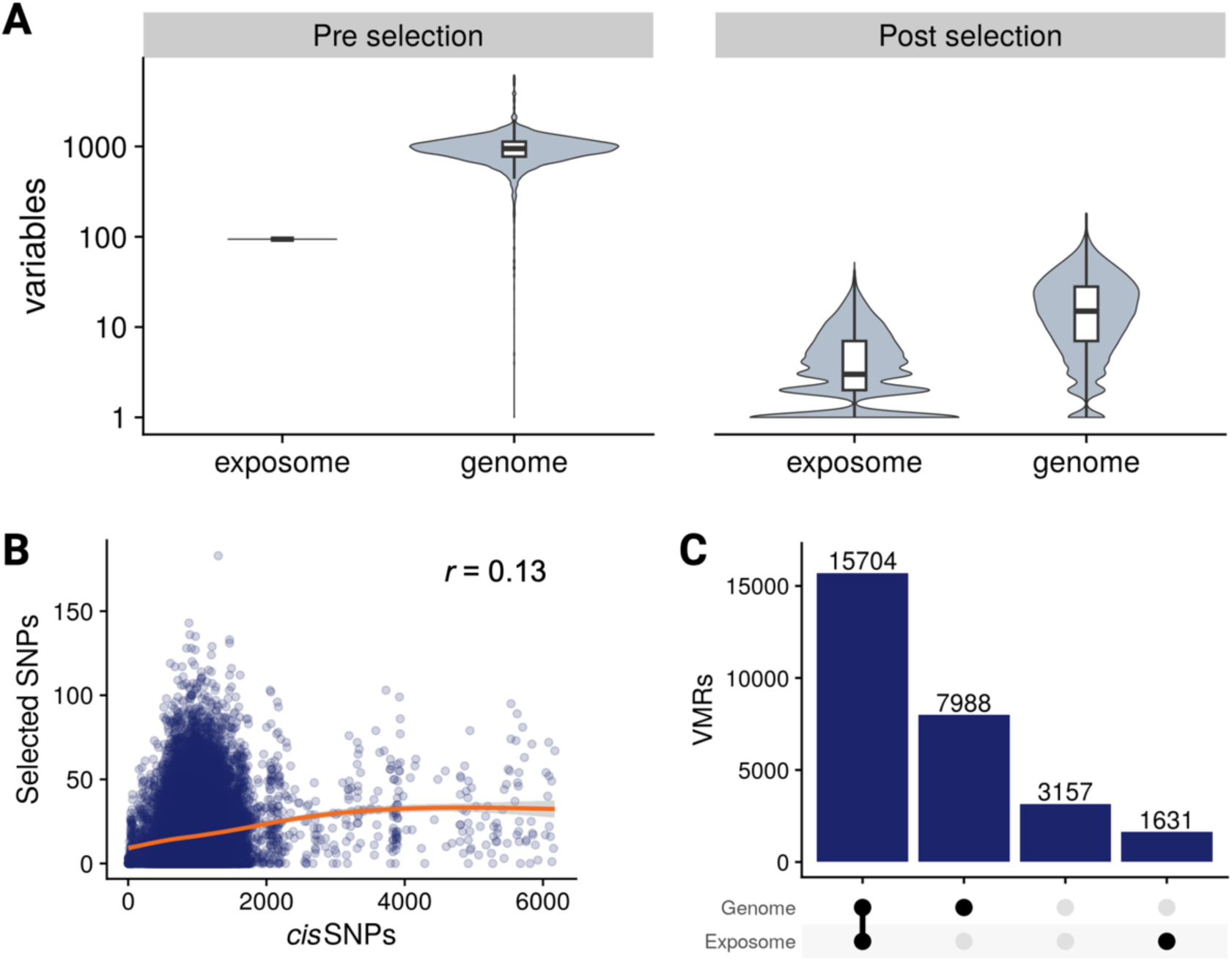
Feature selection reduced and balanced the number of genome and exposome variables per VMR. A) Number of exposome and genome variables in the set of VMRs before (left) and after (right) the feature selection strategy based on LASSO. The initial number of variables for each VMR corresponds to the SNPs in *cis* (genome) and all the 94 available exposome variables. B) Relation of initial and selected SNPs across the VMRs. C) From left to right: number of VMRs with at least one variable selected in both the genome and exposome, only the genome, none, or only the exposome.

Despite reducing the initial difference in the number of genome and exposome variables, we still found a higher number of selected G features compared to E across the VMRs. To explore if this could be an artifact driven largely by genetics having a higher number of variables (initial mean G = 980, initial mean E = 94), which could lead to more selected features by chance, we analyzed the number of selected G and E features in a permuted scenario. In a permuted scenario, any association between DNAme and G or E is broken through data shuffling; after the permutation, differences in the observed number of selected features between groups can be assumed to be driven by the variable imbalance, isolated from the effect of true underlying associations. We observed an overall similar distribution of the number of selected variables, with a median of 0 as expected for a permutated scenario, but a heavier tail for the genome compared to the exposome (genome: mean = 3.7 SPNs per VMR, *SD* = 6.9; exposome: mean = 2.2 variables per VMR, *SD* = 4.2; **Figure S1A**). These results showed that the initial difference in the number of variables exclusively has a minimal overall effect on the number of selected genome and exposome features. The distribution of the number of selected features was highly similar between G and E when we equaled the initial number of genome and exposome variables via random subsampling of G prior to the selection (**Figure S1B**).

When characterizing the selected features, we found a significant enrichment of selected G variables (i.e. SNPs) for *cis* mQTLs (*p* < 0.05, OR = 1.86). We found no relation between the initial and post-selection number of G variables in the VMRs (r = 0.13; **Figure 3B**); this characteristic is especially important to prevent the potential bias towards G models winning in VMRs in densely genotyped regions at the modelling stage, and further supports the robustness of the feature selection strategy to the initial number of variables. Compared to the baseline data set, selected features were not enriched in any prenatal exposome dimension (post selection: maternal health = 51.6%, maternal nutrition = 27.5%, maternal psychosocial status = 9.9%, built environment = 11%). Comparing canonical and non-canonical VMRs, we observed a small difference in the initial number of *cis* SNPs (canonical mean = 1023.9, non-canonical mean = 967.5; *p* < 0.05). After the variable selection, the difference of SNP variables persisted (canonical *mean =* 22.1, non-canonical *mean =* 14.6; *p* < 0.05). Similarly, we observed a small difference in the number of environmental variables post-selection between the VMR categories (canonical *mean =* 3.5, non-canonical *mean =* 2.9; *p* < 0.05; **Figure S2A**).

By the end of the variable selection, 15,704 (55%) VMRs had both E and G variables selected, 7,988 (28%) only G, 1,631 (6%) only E, and 3,157 (11%) neither E nor G (**Figure 3C**). These proportions were similar between canonical and non-canonical VMRs (**Figure S2B**). VMRs with no G and E selected variables occurred when no genome or exposome variables substantially improved the model performance compared to having only the concomitant variables. Because of this, VMRs with no G and E variables selected (11%) were labeled as having Non-Conclusive (NC) results in our data set.

### Genetic variants were included in the majority of VMR best models and explained the largest proportion of DNAme variance

After screening and selecting the potentially relevant variables through a data-driven approach, we evaluated and selected the best model for each of the VMRs with G or E variables selected (89% of them). The evaluated models included one G and/or one E variable at a time to reduce the statistical complexity and computational time involved in assessing multivariate GxE interactions. We fitted all the possible G, E, and pairwise G+E and GxE models using the variables obtained in the feature selection step. For each VMR, the model with the lowest AIC, a penalized likelihood metric, was identified as the best fit. We then implemented a permutation-based strategy to simulate a null distribution and identify VMRs which best model performed no better than expected by chance; these VMRs (30.1%) were labelled as NC in addition to the VMRs that had no relevant G or E variables identified earlier in the feature selection step (11%; total = 41.1%). Out of the informative models (58.9%; i.e., not NC), the G+E model best explained most of the VMRs (27.1%), followed by G (18.3%), GxE (13.4%) and an almost negligible fraction of E (0.1%; **Figure 4A**). This order was consistent between canonical and non-canonical VMRs, with the difference of non-canonical VMRs having a higher proportion of non-conclusive models (46%) compared to canonical regions (24%; **Figure S3A**). In almost all cases (> 99.99%) where our selection strategy picked both G and E variables, the best model in the AIC comparison was either additive (G+E) or interaction (GxE; **Figure S3B**).

**Figure 4.**
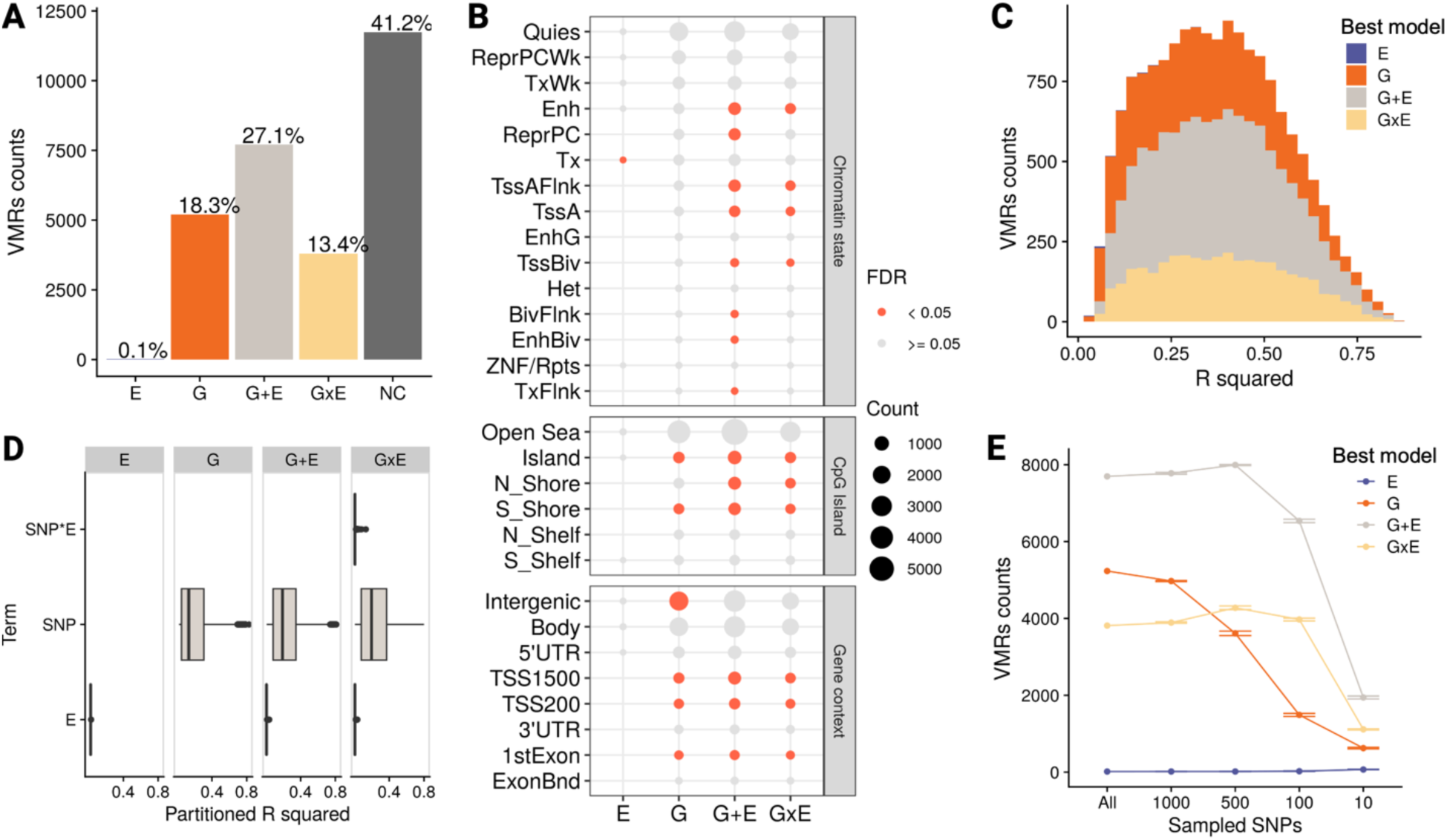
Genetic factors were included in most VMR models and explained the largest proportion of DNAme variance. A) Best explanatory models for the set of VMRs in CHILD cord blood samples. *NC* = Non-Conclusive B) Characterization of model groups regarding CpG island context, chromatin state and gene annotation. Definition of categories: Shore = 0.2 kb from island, Shelf = 2-4 kb from island, N = upstream(5’) of CpG island, S= downstream (3’) of CpG island; TssA = active transcriptional start site (TSS); TssAFlnk = flanking active TSS; TxFlnk = transcribed state at the end of the 5’ and 3’ genes; Tx = strong transcription; TxWk = weak transcription; EnhG = genic enhancers; Enh = enhancers; ZNF/Rpts = ZNF genes and repeats; Het = heterochromatin; TssBiv = bivalent/poised TSS; BivFlnk = flanking bivalent TSS/Enh; EnhBiv = bivalent enhancer; ReprPC = repressed PolyComb; ReprPCWk = weak repressed PolyComb; Quies = quiescent/low; TSS200 = 0–200 bases upstream of the TSS; TSS1500 = 200–1500 bases upstream of the TSS; 5’UTR = Within the 5’ untranslated region, between the TSS and the ATG start site; Body = Between the ATG and stop codon, irrespective of the presence of introns, exons, TSS, or promoters; 3’UTR = between the stop codon and poly A signal. C) R^2^ distribution of best explanatory models that passed the permutation analysis threshold. D) Partial R^2^ of the genetics, environmental and interaction terms estimated with the LMG method. E) Results sensitivity to information reduction in the genome component estimated through the random sampling of SNPs (detailed information in methodology). Error bars represent *SD* (n = 5). The number of NC models were not plotted to improve the readability of the graphic, but this can be obtained by subtracting the number of G, E, G+E and GxE models from the number of total VMRs (28,480) in each under-sampled experiment.

After identifying the informative best models, we characterized the selected variables and VMRs within model groups to unpack specific contributions and understand the underlying fabric of each group. We found an enrichment of mQTLs in the set of SNPs from winning models with a genetic component (i.e., G, G+E and GxE) compared to the set of selected *cis* SNPs (*p* < 0.05, OR = 5.2). The exposure dimension with the highest proportion of variables selected by informative models was maternal health (49.5%), followed by maternal nutrition (28.1%), built environment (11.7%) and maternal psychosocial state (10.7%). This trend was consistent in canonical and non-canonical VMRs (**Figure S4**), and recapitulated the pattern observed in the number of variables selected. We found distinct enriched genomic and functional categories for each model group. G winning-models were enriched in categories related to active Transcription Start Sites (TSSs), G+E in active and bivalent TSSs and enhancers, GxE in active TSSs and enhancers, and E in strong transcription despite the small number of VMRs (**Figure 4B**).

To have a more granular understanding of the contribution of genetic and environmental factors to DNAme, we dissected the explained variance in our results. Informative models explained on average 37% of DNAme regional variance (mean *R*^2^ = 0.37, *SD* = 0.18) with a similar range across model groups except for E, which tended to have smaller values (**Figure 4C**). When decomposing the variance of each VMR to estimate the contribution of each term across the best models, we found genetic terms (mean partitioned *R*^2^ = 0.22, *SD* = 0.18) to consistently explain a significantly higher amount of variance compared to the environmental (mean partitioned *R*^2^ = 0.01, *SD* = 0.004; *p* < 0.05) and interaction (mean partitioned *R*^2^ = 0.01, *SD* = 0.01; *p* < 0.05) terms (**Figure 4D**; full results in **Supplementary Data 2**). This result was again consistent across canonical and non-canonical VMRs (**Figure S5**).

An important consideration when comparing the contribution of the genome and the exposome to DNAme variability is that capturing varying environmental exposures in a population is methodologically more challenging than measuring genetic variants. To explore the sensitivity of our results to this information imbalance, we simulated scenarios with a varying degree of “captured” information of genetic variants by reducing the number of available *cis* SNPs per VMR by random under-sampling. When repeating the RAMEN analysis in this context, we observed an inverse relationship between the number of initial SNPs in the analysis and the VMRs with NC models. Specifically, individual models of the under-sampled factor (G models) were especially sensitive to the information loss, while joint models (G+E and GxE) remained relatively stable (**Figure 4E**). Notably, the number of VMRs best explained only by the unchanged factor (E models) was robust to the different degrees of information in G. These results showed how datasets with a limited characterization of the genome or exposome are likely to underestimate the contribution of their respective genetic or environmental components, and that our method prevents this under-characterization from causing an overestimation of the other component’s contribution.

### Genome-exposome contribution trends were replicated in the PREDO cohort with a complete agreement in the involvement of genetics

Next, we wanted to test whether our genome-exposome contribution results from CHILD replicated in another cohort. To do this, we used RAMEN to reanalyze data from the PREDO study, which was previously investigated in the context of understanding genetic and environment contributions to DNAme in cord blood in 2019 (referred here as C_2019)^23^. Briefly, the PREDO data was composed of two cord blood sets that were profiled for DNAme with the Illumina 450k (PREDO I; n =817) or EPIC array (PREDO II; n = 146). The PREDO data sets captured 8 prenatal environmental variables from the maternal psychosocial and health dimensions (two and six respectively; 6/8 also measured in CHILD). Beyond an attempt at replication, using PREDO allowed us to compare RAMEN with one of the methodologies that we built on. Despite being conceptually similar to C_2019’s approach, RAMEN presents substantial differences as it takes into account the microarray probe design and pairwise CpG correlations to identify VMRs, and implements machine learning and permutation techniques to robustly identify winning models.

Using RAMEN, we identified a lower number of VMRs in the PREDO data sets compared to CHILD, although the VMR number and composition had higher similarity between data sets derived from the same array platform (i.e., EPIC in CHILD and PREDO II; **Table 1**). Out of the informative models in PREDO I, G was the most common winning model, followed by G+E, GxE and E (**Supplementary Data 3**). In PREDO II, the rank order was G, GxE, G+E, and E (**Figure 5A**; **Table 1**; **Supplementary Data 4**). Additionally, we observed a higher proportion of NC models in both PREDO data sets compared to CHILD, and a consistent proportion of G and E models across the PREDO and CHILD data sets (**Table 1**). When dissecting the DNAme variance explained by each term in the winning models, as observed in CHILD, we found the SNP term (PREDO I: *mean =* 0.08, PREDO II: *mean =* 0.16) to explain a significantly higher amount of variance (p < 0.05) compared to the interaction term (PREDO I: *mean =* 0.02, PREDO II: *mean =* 0.06) and environmental term (PREDO I: *mean =* 0.01, PREDO II: *mean =* 0.03; **Figure 5B**).

**Figure 5.**
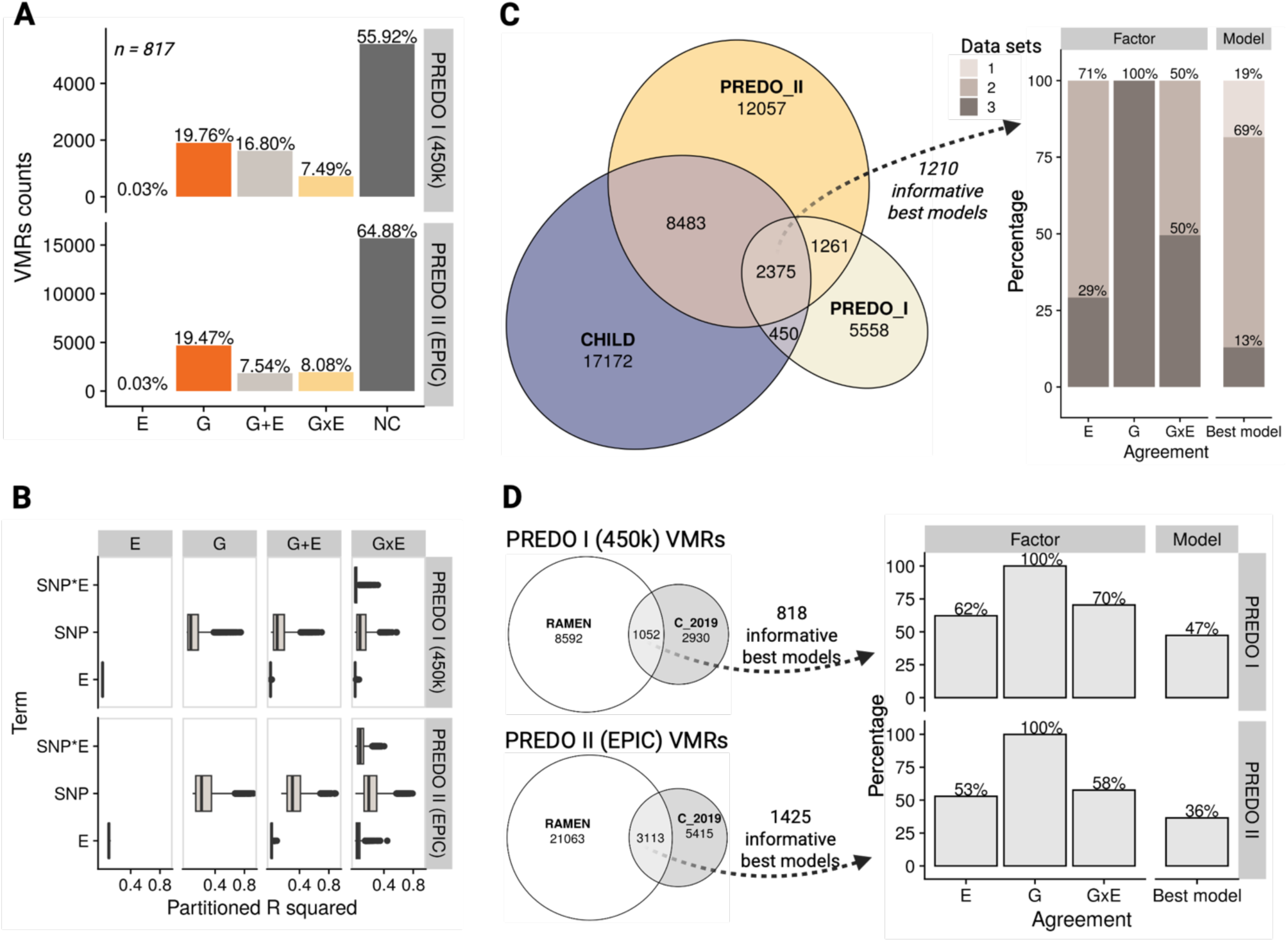
Gene-environment contributions to DNA methylation variation in PREDO recapitulated CHILD results. A) Best explanatory models for the set of RAMEN VMRs in the PREDO cord blood data sets. *NC* = non-conclusive. B) Partial R^2^ of the genetics, environmental and interaction terms. C) Factor and model agreement in overlapping VMRs between the RAMEN and C_2019 methodology in the PREDO data sets. White fill in plots corresponds to results obtained with RAMEN. D) Factor and model agreement in overlapping VMRs between the PREDO I and PREDO II cohorts in the RAMEN and C_2019 results. In the C_2019 comparison all models in the overlap are compared because there is not a non-conclusive category. D) Factor and model agreement in overlapping VMRs between the CHILD, PREDO I and PREDO II data sets using RAMEN. Note that, in the factor agreement, since we were working with 3 data sets and we were comparing a binary state (absent/present), the minimum agreeing data sets that can exist is 2, not 1.

**Table 1.** Results of the genome-exposome contribution to methylome analysis in the CHILD and PREDO studies using RAMEN.

| Cohort | n | VMRs | E models | G models | G+E models | GxE models | NC models |
| --- | --- | --- | --- | --- | --- | --- | --- |
| **CHILD**<br>(EPIC) | 699 | 28,480<br>(6,489 canonical;<br>21,991 non-canonical) | 18<br>(0.06%) | 5,203<br>(18.3%) | 7,710<br>(27.1%) | 3,808<br>(13.4%) | 11,741<br>(41.2%) |
| **PREDO I**<br>(450k) | 817 | 9,644<br>(4,464 canonical;<br>5,180 non-canonical) | 3<br>(0.03%) | 1,906<br>(19.76%) | 1,620<br>(16.8%) | 722<br>(7.49%) | 5,393<br>(55.92%) |
| **PREDO II**<br>(EPIC) | 146 | 24,176<br>(6,598 canonical;<br>17,578 non-canonical) | 8<br>(0.03%) | 4,706<br>(19.47%) | 1,824<br>(7.54%) | 1,953<br>(8.08%) | 15,685<br>(64.88%) |

To compare the similarity between the results obtained in the PREDO and CHILD data sets, we evaluated their agreement. We assessed agreement using two approaches: 1) factor involvement, where we considered a VMR to be consistent for each individual factor (E, G or GxE) if said factor was both absent or both present in the best model, and 2) overall model, where we considered a VMR to be consistent if the best model matched in all factors. For example, if the best model of an overlapping VMR was E in the first set of results (model: sVMR ∼ E + covars) and G+E in the second (model: sVMR ∼ E + G + covars), said VMR would be labelled as agreeing in the E factor (since the E term is present in both models), agreeing regarding the GxE factor (since an interaction term is not included in the winning models), not agreeing regarding the G factor (since the G term is not involved in the first set of results but included in the second), and not agreeing in the overall model (since in the first set of results the model is E and in the second is G+E).

We identified 2375 shared VMRs in the three data sets. Out of the 1210 informative best models (i.e. not belonging to the non-conclusive category in any data set), all of them were in full agreement regarding the G involvement (100%), and less so in GxE (50%) and E (29%). Additionally, 82% of the overlapping VMRs had a consistent overall model in two to three of the data sets (**Figure 5C**). Compared to the C_2019 results, RAMEN displayed an identical inter-data set agreement metric regarding G involvement (100%), an increase in the GxE (+19%) and overall model agreement (+2%), and a decrease in the E model agreement (−12%; **Figure S6**).

Finally, since the C_2019 study used a conceptually similar methodology compared to RAMEN, we compared the results from both methods using the same data (either PREDO I or II) to evaluate their consistency. When comparing the VMRs obtained with the C_2019 and RAMEN methods, we observed a modest VMR overlap in the PREDO I (1,052; 26%) and PREDO II (3,113; 37%) data sets (**Figure 5D**). To evaluate model agreement, we analyzed only overlapping VMRs with informative models in RAMEN, as the Non-Conclusive category is absent in the C_2019 method.

We found a full agreement between the results of the RAMEN and previous C_2019 method regarding the involvement of genetics (PREDO I: 100%, PREDO II: 100%), a modest agreement in the environmental (PREDO I: 62%, PREDO II: 53%) and interaction involvement (PREDO I: 70%, PREDO II: 58%), and a lower agreement in the overall model (PREDO I: 47%, PREDO II: 36%; **Figure 5B**).

## Discussion

In this study, we analyzed multi-omics data to understand the individual and joint contribution of genetic and prenatal environmental differences to cord blood DNAme variability. We developed RAMEN, a user-friendly FAIR^28^ tool built upon previous methodological approaches in the DNAme field to model regional variability with linear models^22,23,31^, while incorporating new strategies to tailor the extraction of regional DNAme variability information for microarrays, and to improve the computational efficiency and genome-exposome contribution estimates. We analyzed the CHILD and PREDO cohorts and found genetics to be a consistent central contributor to cord blood DNAme variation. Our results furthermore provided a detailed catalogue of genomic regions with highly variable DNAme, along with the estimated contribution of genetics, environmental exposures, and gene-environment interactions to their variability. These findings may inform precision medicine approaches on the potential factors associated with the DNAme variation in CpGs of interest, and improve our understanding of the relation between genetic variants, environmental exposures and early life DNAme variability patterns.

We identified a higher number of VMRs compared to previous perinatal studies even after implementing a within-region probe correlation filter. The number of VMRs found with RAMEN and their type composition suggests that the increment is driven mainly by our implementation of non-canonical VMRs. Non-canonical VMRs consider the sparse and non-homogeneously covered design of the microarray, by far the most used technology to profile DNA methylomes in epidemiological cohorts. Non-canonical VMRs are single probes in the array that were filtered out in previous methodologies solely due to sparsity. Taking these sites into account is especially important in the EPIC v1 array, since the DNAme levels of most regulatory regions are measured targeting single probes that accurately represent the methylation state of the surrounding sites, as reported in empirical comparisons with whole-genome bisulfite sequencing data^32^. Our framework optimized the extraction of variability patterns by considering this design, which led to a 3-fold increase in the number of genomic regions analyzed. Even though the VMR identification criteria was created for the EPIC array, we observed them to be equally informative in the 450k microarray. Additionally, the non-random VMR genomic distribution we observed is consistent with previous studies reporting highly variable DNAme sites being enriched in open sea, intergenic, transcription, enhancer, quiescent and repressed regions^23,33,34^.

A key strength in our study was the computationally efficient detection of non-conclusive (NC) models, which behave like spurious associations expected by chance. With this methodological step, we expect our results to provide a likely smaller, yet more refined estimation of the DNA methylome-wide influence of genetics and the environment. We found genetics as a key contributor in most informative cord blood VMR models, primarily in combination with the environment, whereas the environment alone explained a, notably small proportion of DNA methylation variability. This trend aligned with similar studies in cord blood, umbilical cord and placenta^22–24^; however, our G, E, G+E and GxE proportion estimates differ substantially.

We found 36-59% of methylome-wide VMRs associated with genetics, significantly lower compared to previous reports in perinatal integrative studies estimating jointly the contribution of genetics and the environment (96-100%)^22–24^. Our estimate aligned more closely with studies using significantly different methodological approaches to interrogate specifically the association of genetics with DNAme, such as the largest mQTL blood study to date (44% of CpGs under *cis* genetic control)^35^ and the narrow-sense heritability estimates in twins (41% of CpGs with significant additive genetic effects)^36^. Similarly, our estimated proportion of VMRs with gene-environment interactions on DNAme (∼10%) is lower than those reported in gene-environment integrative studies in umbilical cord (75%)^22^, cord blood (53% on average across 3 cohorts; SD = 8.6)^23^ and placenta (70%)^24^.

VMRs best explained by G, E, G+E and GxE were enriched in distinct locations important for genomic regulation. Consistent with our findings, twin studies in blood at age 18 and epidemiological cohorts at birth in cord blood and placenta have reported DNAme sites with a genetic contribution to be enriched in regions within a 5 kb window upstream of TSSs^37^, and sites with an environmental and genetic contribution (G+E) to be enriched in the immediate vicinity of TSS, the 1^st^ exon of genes, CpG islands, bivalent enhancers and regions repressed by Polycomb^23,24,37^. These findings suggest distinct, potentially conserved pathways through which genetic and environmental factors might associate with variability in the DNA methylome and potentially affect cellular processes across the first 18 years of life.

When dissecting the contribution of each factor in the winning models, we found genetics to explain a higher amount of DNAme variance in cord blood VMRs compared to the environment and interaction terms, and to be fully consistent in overlapping VMRs across the CHILD and PREDO cohorts. The estimated contribution of genetics to DNAme variance across VMRs (*mean =* 22%) was consistent with the average narrow-sense DNAme heritability estimations in blood family studies (i.e., proportion of variance that can be attributed to additive genetic effects), which ranges from 0.1 to 0.3^10,36,38–42^. We also observed linear GxE interactions to have small contributions (mean partitioned R^2^ = 0.01). This suggested that large sample sizes might be required to detect GxE interactions in the DNA methylome, which may also be related to the limited number of significant GxE associations documented in the few DNAme studies looking at epigenome wide GxE^15–19^. VMRs best explained by GxE models in our results may further be used to conduct candidate gene-environment interaction studies, which could help decrease the multiple testing burden and refine precision medicine approaches^20^.

The contribution of environmental factors was less consistent than SNPs in overlapping VMRs across cohorts (i.e., E and GxE terms). This might be a consequence of the E and GxE factors having a lower average contribution to the DNAme variance, requiring a greater statistical power to be consistently detected. One explanation for this low contribution could be the presence of specialized extra-embryonic tissues during gestation, such as the placenta, that protect the fetus and directly regulate the prenatal environment^43,44^. These extra-embryonic tissues might dampen the overall environmental exposures effect in cord blood. Alternatively, the low concordance and environmental contribution might result from known biases affecting the power to detect and replicate exposome effects, such as the small number of measured variables, heterogeneity of measurements and their errors, reliance on indirect exposures assessments (e.g., self-reported questionnaires), difficulty capturing both the effector exposures at the relevant timepoint and its duration, lack of standardized protocols to measure the heterogeneous and dynamic nature of the exposome, low effect sizes, and differences in the exposures measured across epidemiological studies^45–47^.

A significant proportion of VMR were identified with a Non-Conclusive model, which means that none of our G and E variables, individually or jointly, substantially improved the model performance. We observed that this proportion increased with higher rates of missing information, which was evident in the SNP sub-sampling experiment in CHILD and the higher NC proportion in PREDO I compared to CHILD (similar sample sizes but fewer E variables in PREDO I). Additionally, we observed a higher proportion of NC models in PREDO II compared to PREDO I (same number of G and E variables but a smaller sample size in PREDO II). Lastly, the DNAme of some VMRs might be governed by stochastic processes^48^. Therefore, we hypothesize that NC models may be driven by a combination of unmeasured variables, sample size limitations, and stochasticity.

Finally, our study has limitations and challenges that must be considered when interpreting results. Regarding the method, our framework is designed to assess gene-environment interactions only if the corresponding SNP and exposure are potentially relevant individually, since we conduct feature selection before fitting the models. Additionally, in cases of high correlation between individual G or E variables (e.g. groups of SNPs with a large amount of linkage disequilibrium), our feature selection strategy will select only one at random for each VMR, meaning our results should be interpreted as associations at the G or E level. Refined strategies and study designs are needed to fine-map causal gene-environment combinations, which can be particularly challenging in cases of gene-environment correlation (*i.e.,* different genotypes systematically experiencing specific environments)^49^.

To reduce computational time and model complexity, we modeled variation as univariate in G and E, although combinations of multiple G or E factors might contribute to DNAme variability to a smaller degree. Given the polygenic nature of DNAme^35^, approaches incorporating multiple G or E variables and their interactions could improve the variance explanation, although the number of possible models was considered too high to incorporate into our framework. Additionally, our method estimated the variance explained by each G, E and GxE factor in the best informative models after accounting for the variance explained by the concomitant variables; the variance explained by genetic and environmental factors might be underestimated when collinear with the concomitant variables.

Finally, our study provides a comprehensive characterization of gene-environment effects on cord blood DNA methylome variation. Yet, further studies will be helpful to understand this phenomenon in additional contexts. Regarding the platform used to profile the methylome, microarrays are designed to target primarily non-repetitive regions. DNAme at repetitive regions, such as sub-telomeric repeats and rDNA sequences, have been reported to be especially responsive to the environment^50,51^. Long-read sequencing technologies are needed to explore the G and E contribution in these genomic elements. On a temporal matter, our study focuses on the prenatal period. As twin studies have suggested that environmental effects on DNAme variance increase with age in 10.4% of CpGs^36^, longitudinal studies will be instrumental to assess the gene-environment contribution across the human lifespan. Lastly, our cohorts primarily included socio-economically privileged individuals from Western, Educated, Industrial, Rich and Democratic (WEIRD) countries. Studying genetically and/or socioeconomically homogeneous populations can bias the characterization of VMRs and the estimation of environmental and genetic effects. Systemic efforts to discontinue colonialist and unethical practices are needed to include historically underrepresented populations in research, clarify the role of genetics and the environment in the methylome and conduct research relevant to all individuals in our society.

## Conclusions

We identified genetics as a key contributor to DNAme variability in cord blood, frequently in combination with the prenatal environment, supported by a large fraction of informative models including G, a high proportion of DNAme variance explained, and full consistency across cohorts. We reported a detailed genomic map of regions with high interindividual DNAme variation and identified the potential factors explaining their variation, which could be a powerful resource to annotate CpGs and identify GxE interactions in perinatal cohorts. We also introduced a FAIR tool to estimate the genome-exposome additive and interactive contribution to the DNA methylome. Our findings highlight the importance of accounting for genetic effects in DNAme studies, and contribute to the understanding of how and under which conditions individual genetic susceptibility and environmental variables may work together to influence DNAme in a highly sensitive developmental period.

## Methods

### Cohort descriptions

#### CHILD

The CHILD study is a national longitudinal birth cohort following infants from prenatal to age 8 years (with older assessments ongoing) with participants across four provinces in Canada: British Columbia, Alberta, Manitoba, and Ontario. Inclusion and exclusion criteria can be found in the cohort description publication ^25^. For the current study we used a subset of 699 cord blood samples from term (>36 weeks) newborns with genome, methylome and exposome data that passed quality control; this subset of samples within the larger CHILD cohort was selected for multi-omic profiling because of the availability of longitudinal biological samples at birth, age one and age five. The self-reported ethnicity of these newborn’s mothers was 76.7% Caucasian White, 13.2% Asian (including East Asian, South Asian and South East Asian), 3% First Nations, 1.6% Black, 1.1% Hispanic and 4.4% other (including multiethnic and Middle Eastern), and their average age was 33.3 (*SD*=4.6). The assigned sex at birth of the individuals was 46% female and 54% male. Gestational age at delivery was 39.7 weeks on average (*SD* = 1.2 weeks). The distribution of these demographic variables were similar to the ones in the full study^26^.

#### PREDO

The PREDO study is a longitudinal multicenter perinatal cohort from Finland^52^. Briefly, the study recruited pregnant women and their singleton children born alive between 2006 and 2010. Inclusion criteria were either pregnant women with known clinical risk factor status for preeclampsia and intrauterine growth restriction (IUGR), or pregnant women who volunteered to participate who did not present risk factors. Further details about study design can be found in the cohort description publication^52^. For this study, we used a subset of 963 cord blood samples from ethnically homogeneous Finnish newborns with genome, methylome and prenatal exposome data.

### DNA methylation

#### CHILD Sample collection and DNAme profiling

Specimens were collected directly from an umbilical vein before the placenta was delivered in heparinized vacutainers, pooled in a 50 mL conical tube, aliquoted and frozen^26^. DNA was extracted from cord blood samples using the DNeasy Blood & Tissue Kit (Qiagen), samples were bisulfite converted using EZ-96 DNA Methylation kit (Zymo Research) and DNA methylation profiles of the samples were measured with the Infinium MethylationEPIC BeadChip v1 array (Illumina).

##### CHILD DNAme Pre-processing

Cord blood DNAme data was pre-processed and subjected to quality control checks in R v4.0.3. Sample exclusion criteria were the following: failure in *EWAStools* v1.7^53^ control metrics, reported-predicted sex mismatch inferred using sex chromosome intensities (*minfi* v1.44^54^), EPIC-GSA SNP mismatch, maternal cord blood contamination^55^, detection *p* value, >1% missing probes measured and outliers (detected with the *lumi*^56^ package). Normalization was next conducted using BMIQ with noob to account for probe type bias and background correction implemented in the *wateRmelon*^57^ v2.4 and *minfi* v1.44 packages, respectively. Probe exclusion criteria were the following: SNP probes, detection *p* value > 0.01 in > 5% samples, present in Pidsley *et al.*’s cross-hybridizing and polymorphic EPIC probes^32^, used in Haftorn’s 2021 EPIC gestational age clock^58^ or in the *FlowSorted.Blood.EPIC*^59^ cord blood IDOL’s probes, and located in non-somatic chromosomes. As a result of these steps, 785,689 out of 866,836 CpG probes and 813 out of 828 samples were retained. Finally, batch effects associated with technical variation (chip and row) were removed using the ComBat function from the *sva* package^60^.

#### PREDO DNAme

The DNAme data pre-processing was conducted as described previously^23^. Briefly, the data set consisted of 817 cord blood samples that were ran in Illumina 450k Methylation arrays, and 146 in Illumina EPIC arrays. We conducted quality control filters regarding minfi^54^ control metrics, median intensity, reported-estimated sex and maternal DNA contamination, CpG detection p value. Methylation beta-values were normalized using the *funnorm*^61^ function, and variation associated with batch effects was further removed using the ComBat function from the *sva* package^60^. Probes in sex chromosomes and reported to be cross-hybridizing were excluded. The final data set consisted of 418,790 and 794,414 CpG probes for the Illumina 450K and EPIC arrays respectively.

### Genotyping

#### CHILD Pre-processing

DNA was extracted from cord blood samples as mentioned in the DNAme section, and genotyping was performed using the GSA v3 + Psych v1 array (Illumina). Following GenomeStudio v2.0.4 (Illumina) preliminary quality control checks, we exported the genotyping data to R v4.2.0 for further QC guided by Illumina recommended metrics^62^. Detailed thresholds can be found in **Supplementary Methods**. Sample relatedness was checked by calculating kinship coefficients (identity by descent) based on Maximum Likelihood Estimation (MLE) in the R SNPRelate package^63^. Based on a kinship coefficient score of 0.5, 3 pairs of related samples were identified which were technical replicates and being sample duplicates were expected to be identical. Only one sample from the 3 pairs was retained for downstream analysis. Probes in non-autosomic chromosomes were excluded from the analysis. Imputation was conducted using the ENIGMA imputation protocol^64^ based on the Michigan Imputation Server pipeline^65^. After imputation, imputed SNPs with an R^2^ £ 0.8 and MAF £ 0.01 were filtered out using *bcftools* v1.16^66^. Finally, we conducted LD pruning using *plink* 1.9^67^ with a window size of 50 variant counts, a 5 variant count window shift and a 0.5 pairwise R^2^ threshold. The final genotyping consisted of a data set of 1,260,703 SNPs (492,062 directly measured) and 824 out of 826 individual samples.

#### PREDO genotyping pre-processing

The genotype data pre-processing was conducted as described previously^23^. Briefly, genotyping was performed on Illumina Human Omni Express Exome Arrays containing 964,193 SNPs. We conducted quality control filters regarding call rate, MAF, HWE, relatedness, reported-predicted sex and heterozygosity outliers. Following imputation, we ran another round of quality control regarding SNP info score, MAF, HWE, *p* value and call rate. Next, the genotype data set was pruned using a threshold of r2 of 0.2 and a window-size of 50 SNPs with an overlap of 5 SNPs. The final data set consisted of 983 samples with 788,156 SNPs.

### Prenatal exposome

#### CHILD prenatal exposome

We used 94 prenatal environmental variables collected in the CHILD cohort encompassing four fetal exposure dimensions: maternal psychosocial state, maternal nutrition, maternal health and built environment. Briefly, the maternal health dimension included 46 variables such as conditions during pregnancy (e.g. cold, diabetes, allergy, hypertension, etc.) and smoking; the maternal nutrition dimension included 29 variables such as dietary patterns, Healthy Eating Index 2010 scores, and dietary constituents that mediate 1-carbon metabolism; the parental psychosocial dimension included 10 variables such as maternal depressive symptoms and socio economic status; and the built environment dimension included 9 variables such as NO_2_, PM_2.5_ and O_3_. Detailed information about the pre-processing of these variables can be found in **Supplementary methods**.

#### PREDO prenatal exposome

Of the PREDO data set collected, we included 10 prenatal environmental variables described previously^23^: maternal depressive and anxiety symptoms, maternal treatment with betamethasone, delivery mode, parity, maternal age at delivery, pre-pregnancy BMI, hypertensive pregnancy disorders (including gestational hypertension, chronic hypertension and preeclampsia), gestational diabetes and glucose values during a 75g 2 hour oral glucose tolerance test (OGTT).

#### Prenatal exposome pre-processing

For CHILD, each dimension was pre-processed independently but using the same criteria as described: 1) Individuals with > 30% of missing values were removed; 2) Variables that displayed > 15% of missingness were removed; 3) Variables that measured a similar object and were highly correlated (*r* > 0.8) or displayed low variability (< 10 cases in binary variables) in the dataset were removed; 4) Remaining missing values were single imputed using the *mice* package in R^68^ with 100 iterations and the default method. For the psychosocial dimension, household income when the child was 1 year old was included as a covariate in the imputation due to its high correlation with income at birth and therefore its potential to improve the imputation accuracy. After this pre-processing, the four dimensions were combined into a single dataset, and individuals with missing data in one or more dimensions were removed, which resulted in a dataset of 699 out of 790 individuals with complete environmental exposure information of 94 variables (maternal nutrition = 29, maternal health = 44, maternal psychosocial = 10, built environment = 9). A full list of the variables included for this study and their descriptive statistics can be found in **Supplementary Data 1**.

The PREDO data set was pre-processed in the same way treating all the variables as a single dimension. The final data set consisted of 8 environmental variables: all the mentioned in the PREDO prenatal exposome section except for betamethasone and OGTT.

### Concomitant variables

#### CHILD

The following DNAme confounders, which can lead to biased or spurious results, were considered “concomitant” variables: predicted cell type composition and population stratification (captured by global genetic ancestry). Additionally, the following sources of DNAme variation not directly related to genetic or environmental variation were also considered concomitant: sex (Female/Male) and gestational age (days at delivery). Concomitant variables were included as covariates in all models of DNAme variability

Cord blood cell type proportions of CD8+ T cells, CD4+ T cells, Natural Killer, B cells, Monocytes, Granulocytes and nucleated red blood cells were estimated from raw DNAme data using the *FlowSorted.Blood.EPIC* R package^59^ with the Noob pre-processing method and both the IDOL and hypo/hypermethylated probes to distinguish cell types. To address the compositional and multicollinear nature of cell type proportions, we applied an isometric logratio transformation followed by a robust PCA as proposed by Filzmoser et al. in 2009^69^. The first four components’ eigenvectors were used, which explained 92% of the variance.

Global genetic ancestry was estimated by projecting each sample’s genotypes on the principal components calculated using the reference 1000 Genomes Project phase 3 data^70^. The eigenvectors of the first four principal components, which captured the population stratification by visual assessment, were used.

#### PREDO

Like CHILD, the newborn’s sex (F/M), predicted cord blood cell type proportions, genetic ancestry and gestational age at birth (days) were used as covariates in all models. To compare our results with the ones obtained previously using this cohort, the variables were obtained as described before^23^. Briefly, cord blood cell counts were estimated and incorporated directly into the models (all but one to address their compositional nature). The eigenvectors of the first two genetic PCs were used to capture genetic ancestry.

### Genome-exposome contribution to methylome variability analysis

The genome-exposome contribution to methylome variability analysis was conducted with the RAMEN v1.0.0 R package (**Figure 1**). The sections below detail its methodological steps.

#### Identification of Highly Variable Probes

To identify Highly Variable Probes (HVPs) without the effect of known DNAme confounders, we first regressed out the linear effect of the concomitant variables (sex, gestational age at delivery, cell type proportions and genetic ancestry) on DNAme M values (**Figure 1A, Step 1.1**). The residuals of this regression were used to compute the variance of each measured CpG probe (**Figure 1A, Step 1.2**), and a cutoff of the 90^th^ percentile of the distribution was used to identify HVPs (**Figure 1A, Step 1.3**). Residual M values were only used for HVP identification; unadjusted M values were used in the rest of the analysis.

#### Identification of Variable Methylated Regions

HVPs were grouped into Variable Methylated Regions (VMRs; **Figure 1A, Step 2.1**). We divided VMRs into two categories: canonical and non-canonical. Canonical VMRs are regions that meet traditional grouping criteria: two or more HVPs that are close together (chained in <1kb windows) and share a similar DNAme profile (*r* > 0.15 based on Gatev *et al.*’s^71^ simulation to empirically determine DNAme regional correlation thresholds). On the other hand, we created the non-canonical VMR category to account for the sparsity and non-homogeneous coverage in the EPIC microarray design. This array has a high number of DNA regulatory regions measured by single probes that accurately represent regional DNAme levels^32^. With this category, we aimed to recover valuable information from HVPs that had been excluded in previous studies solely because of the genomic array design. DNAme level of each region was further summarized per individual by taking the median DNAme value per region (**Figure 1A, Step 2.2**).

#### Genome and exposome variable selection

For each VMR we pre-selected their *cis* SNPs (within a 1 Mb upstream and downstream window; **Figure 1A, Step 2.3**). Then, we implemented a scalable variable-selection strategy prior to the model fitting and comparison to reduce the computational burden and address the substantial imbalance in the number of genome and exposome variables, which can introduce a methodological model selection bias towards the category of variables with a substantially higher number of variables. We conducted the variable selection to keep the potentially relevant genome and exposome variables using a method based on Least Absolute Shrinkage and Selection Operator (LASSO; **Figure 1A, Step 3.1**). LASSO is an algorithm with a variable screening property that penalizes more complex models (i.e., with more variables) in favor of simpler ones (i.e., with less variables), but not at the expense of reducing the predictive power. This decreases the number of variables in a model, the downstream computational time and can improve model accuracy^72^. Briefly, for each VMR we conducted a LASSO regression in three scenarios: 1) in the presence of only the *cis* SNPs, 2) in the presence of only environmental variables, and 3) in the presence of both *cis* SNPs and environmental variables. Concomitant variables were included in all models and were unpenalized during LASSO. Further details on the implemented LASSO approach can be found in **Supplementary methods**. For each VMR, all variables selected in scenarios 1-3 (*i.e.,* with a coefficient > 0) were pooled and moved to the model fitting stage.

#### Selection of best explanatory model

For each VMR, all selected genome (SNP_i_) and exposome (E_j_) variables were individually used to fit the models in **Table 2** (**Figure 1A, Step 3.2**). Following the model fitting, the best explanatory model for each VMR was selected based on the models’ lower Akaike Information Content (AIC; **Figure 1A, Step 3.3**). The use of this metric assumed that most of our models were a simplification of the underlying truth (where more than one G and E variables are likely to contribute to the methylome variance). Briefly, AIC penalizes models with higher number of terms, and excels in problems where all models in the model space are considered incorrect and in cases where the selected model could belong to a very large class of functions describing complex relations^73^. Following the model selection, the variance explained by the SNP and E terms in the best models was estimated using the *relaimpo* R package^74^ which uses the Linda, Merenda and Gold (LMG) method^75^.

**Table 2.**
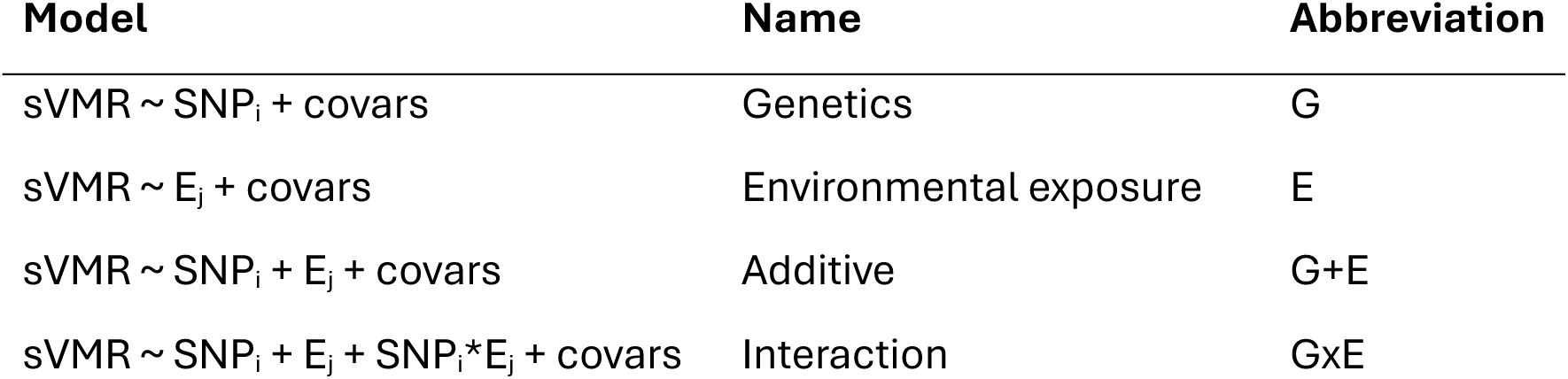
Models fitted and compared in RAMEN per VMR. Where *sVMR* represents the summarized regional DNAme level, *SNP*E* the gene-environment interaction term, and *covars* the previously described concomitant variables (sex, gestational age at delivery, cell type proportions and genetic ancestry).

Finally, to filter out best explanatory models which G/E contribution could be explained by chance, we conducted a permutation analysis (**Figure 1A, Step 3.4**) of the G and E contribution to VMRs’ DNAme. To do so, we shuffled the G and E variables together in the data set and repeated the evaluation of G and E contribution (**Figure 1A, Steps 3.1-3.3**) 10 times. We used the R^2^ G/E increment (*i.e.,* R^2^_BEM_ – R^2^_BM_; where *BEM* = Best Explanatory Model and *BM* = Basal Model including only the concomitant variables) of those permutations to create a null R^2^ increment distribution. We obtained a bimodal R^2^ G/E increment distribution that we stratified into two groups: marginal (G and E), and joint (G+E and GxE) models. The 95^th^ percentile of the null distributions was used as a cut-oi to remove poor performing best models in their corresponding strata.

### Total number of G, E, G+E and GxE models without variable selection

We computed the total number of models that would have been fitted and compared without the variable selection stage as follows:

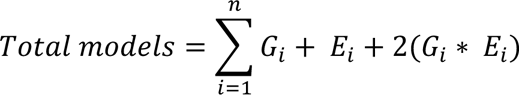

Where *n* is the total number of VMRs and *i* its index, *G* is the number of *cis* SNPs in a given VMR, and *E* is the number of environmental exposures (94 for all VMRs). In lay terms, for each VMR, the number of models fitted is the sum of a) one model for each SNP in *cis*, b) one model for each environmental exposure (E = 94), and c) two models (one additive and one interaction) for each possible SNP and E combination. The sum of this was 5,279,537,086 models.

### Comparison of HVPs with Derakhshan’s highly variable CpGs

To compare the HVPs in our study with previously reported human tissue- and ethnicity-independent highly variable CpGs (hvCpGs)^33^, we downloaded the list of probes from the supplementary data in Derakhshan *et al*.’s publication. In summary, they analyzed 30 data sets that covered 8 ethnicities and 19 tissues/cell types spanning a wide range of ages (0.75-108 years old). There, hvCpGs were defined as probes covered in at least 15 of the 30 data sets and with a methylation variance in the top 5% of all (non-removed) CpGs in ≥ 65% of data sets where the CpG was covered (after qc). Since this hvCpGs data set was made with Illumina 450k array data, we conducted the comparison only in the subset of probes present in both the EPIC and the 450k array.

### mQTL analysis

We conducted an mQTL analysis using the R package *Matrix eQTL* v2.3^76^ on a superset of 790 cord blood samples profiled for DNAme and genotyped as described above. These 790 samples correspond to the 699 individuals used for the genome-exposome contribution to methylome analysis plus 91 samples from the same cohort (CHILD) that were excluded from said analysis due to incomplete exposome information. The covariates included in the association test are the same ones described in the *Concomitant variables* section above. *Cis* (SNP-CpG pars within 1 Mb) mQTLs were detected with a p-value threshold of 1×10^−5^. The significant *cis* mQTLs found in the 207,199 CpGs (26.47% of the probes) can be found in **Supplementary Data 5**.

### Enrichment analyses

#### Enrichment of mQTLs in selected and best-model SNPs

To test if A) the SNPs selected in the variable selection step and B) the SNPs present in the best model per VMR (if G was a component of the selected model) were more likely to be mQTLs, we conducted a chi-squared test. For the over enrichment analysis in A we used the selected *cis* SNPs as target and all the SNPs in *cis* of a VMR as background. For the enrichment analysis in B, we used the *cis* SNPs in all the winning models as target and the selected *cis* SNPs as background

#### Enrichment of CpG categories in VMRs and best models

We used the CpG manifest provided by Illumina to annotate the CpG probes regarding gene annotation and island context using the *UCSC_RefGene_Group* and *Relation_to_UCSC_CpG_Island* columns. Additionally, we used the core 15-state model of cord blood T cells (epigenome E033) generated by the Roadmap epigenomics project^77^ (downloaded from https://egg2.wustl.edu/roadmap/web_portal/chr_state_learning.html) to annotate the CpG probes regarding chromatin state. We then conducted an over-representation analysis of each CpG category by doing a one side Fisher’s test with the *clusterProfiler* R package^78^. For the over representation analysis in VMRs we used all the probes in VMRs as target and the probes in the EPIC array as background. For the over representation analysis in best models, we annotated each VMR according to its probes

(*e.g.*, if a VMR was composed of two probes that were annotated to Tx and Enh respectively, we annotated said VMR as Tx and Enh); we then used the VMRs in each best model group as targets and all VMRs as background.

### Information imbalance analysis

To explore the effect of the genome-exposome information imbalance in the contribution to DNA methylome variability analysis, from the set of *cis* SNPs we randomly sampled 10, 100, 500 and 1000 SNPs for each VMR. In the cases where a given VMR had less *cis* SNPs than the required number to be sampled, no random sampling was conducted. Next, the entire G&E contribution modeling phase of the analysis was conducted (**Figure 1A3**) to get the proportion of best models in each model category. This procedure was repeated 5 times to estimate the variability of the results.

### Variable imbalance analysis

To dissect the effect of the genome-exposome variable imbalance in the contribution to DNA methylome variability analysis, we chose to explore this in a null hypothesis scenario. In a context where the genome and exposome have no effect on DNAme variability (*i.e.* no information imbalance in the variables since none of them are associated to DNAme), we could expect the variable number imbalance to be the main driver of the differences in G and E best models’ proportion, therefore exploring the sensitivity of the method itself. We shuffled the G and E variables together in the data set and conducted the G and E contribution part of the analysis (**Figure 1A3.1-3.3**) with the original number of *cis* SNPs (*i.e.,* a higher amount of G variables than E).

### Model and factor agreement

Overlapping VMRs were defined as regions with at least one overlapping genomic position and were identified with the *plyranges* R package v 1.18.0^79^. Model agreement in overlapping VMRs was evaluated in two modalities: A) overall model and B) factors. To improve comparability, non-conclusive (NC) models were excluded from all comparisons, since NC models can increase the non-agreement percentages due to confounding factors such as differences in information content between cohorts, sample size, and methodology (specifically regarding Czamara-2019^23^ where the NC category does not exist). For (A), we treated each best model as an independent category and labelled overlapping VMRs from two datasets as concordant if they belonged to the same group (e.g., best model being G+E in both data sets). To account for the fact that models are not independent categories, but rather reduced forms of a full interaction model, we next evaluated the individual factor agreement. For (B), overlapping VMRs from two datasets were labelled as agreeing regarding G, E or GxE if said factor was both present or both absent in the best model. For example, if one model were E (model: sVMR ∼ E + covars) and the other G+E (sVMR ∼ E + G + covars), that VMR would be labelled as agreeing regarding E, since E is present in both models, agreeing regarding GxE, since the model agrees that an interaction does not best explain the DNAme, and not agreeing regarding G since G is not involved in the first scenario but included in the second.

## Supporting information

Supplementary data description

## Acknowledgments

We thank the families that took part in the CHILD and PREDO cohort studies for their dedication and commitment to advancing health research, as well as the team involved in the study design, sample collection and data availability, such as interviewers, and clinical and administration staff. We are grateful with Dr. Angela Devlin and Dr. Alejandra Wiedeman for providing valuable input for processing the nutrition variables, Dr. Meingold Chan for providing guidance in the processing and scoring of the psychological questionnaires Dorothy Lin and Carlos Cortés-Quiñones for helping with the creation of the RAMEN logo, and Marcia Jude for conducting the global genetic ancestry analysis in the CHILD samples. This work was supported by BC Children’s Hospital Research Institute Establishment Award (to K.K.). E.I.N.D. was funded by the Graduate Globalink Fellowship from MITACS, the Society to Cell Clyde Hertzman Memorial Fellowship from the Social Exposome Cluster, the Gertrude Langridge Graduate Scholarship in Medical Sciences, the Patrick David Campbell Graduate Fellowship, the Bank of Montreal Graduate Fellowship and the 4-Year PhD Fellowship from the University of British Columbia. The CHILD Cohort study was supported by core funding from The Allergy, Genes and Environment Network of Centres of Excellence (AllerGen NCE) and the Canadian Institutes of Health Research (CIHR); for further information visit childstudy.ca. CHILD epigenetic analysis at the University of British Columbia was funded by CIHR, AllerGen NCE and Genome Canada and Genome British Columbia ([274CHI]). S.E.T. holds a Tier 1 Canada Research Chair in Pediatric Precision Health and the Aubrey J. Tingle Professor of Pediatric Immunology. M.S.K. is the Canada Research Chair in Social Epigenetics and the Edwin S.H. Leong Chair in Healthy Aging - a UBC President’s Excellence Chair. The PREDO study was supported by funding from the Academy of Finland, European Union’s Horizon Europe research and innovation program under grant agreement No 101057390 (HappyMums), Helsinki Institute of Life Sciences (HiLife) Fellows Programme 2023-2024, and HUS VTR (A special Finnish state subsidy for health science research).

## Author contributions

E.I.N.D., K.K. and M.S.K. conceptualized the project. E.I.N.D. conceived the analyses, developed the RAMEN package, conducted the exposome pre-processing, genotype imputation and genome-exposome contribution analysis in CHILD and wrote the manuscript. D.C. conducted the data pre-processing and genome-exposome contribution analysis using RAMEN in PREDO. K.E. conducted the CHILD DNAme pre-processing. E.I.N.D and K.E. conducted the CHILD genotype pre-processing. E.I.N.D, K.K, M.S.K., M.P.F., S.M.M., and C.K. made intellectual contributions to the project analysis and results interpretation. J.L.M. and D.T.S.L. ran the DNAme and genotyping arrays in CHILD. K.K. supervised the statistical analyses. P.M, T.J.M, E.S., P.S. and S.E.T. conceptualized and planned the CHILD study and data collection. K.R. conceptualized and planned the PREDO study and data collection. K.R. and J.L. were involved in the cord blood DNAme profiling in the PREDO study. All authors contributed to and approved the final version of the manuscript.

## Data availability

The participant data is not publicly available for the protection of participants of the CHILD and PREDO studies. Access to this data can be obtained by request; if interested, contact the PREDO **Error! Hyperlink reference not valid.** or CHILD study boards. Summary statistics and results are available in **Supplementary Data 1-4**.

## Code availability

The code used for the analysis of this paper is available in supp. The RAMEN package, which contains the reproducible pipeline for the genome-exposome contribution to DNAme variability analysis, can be found in https://github.com/ErickNavarroD/RAMEN. A full documentation and tutorial of the package can be found in https://ericknavarrod.github.io/RAMEN/articles/RAMEN.html. The code used to conduct the analyses in this manuscript can be found in https://github.com/ErickNavarroD/NavarroDelgado_2025_VMRs.

