## Supplementary data description for "RAMEN: Dissecting individual, additive and interactive gene-environment contributions to DNA methylome variability in cord blood"

**Supplementary data 1.** Statistics and details of CHILD exposome variables.

**Supplementary data 2.** Results of the genome-exposome contribution analysis to methylome variability in CHILD cord blood samples. The file contains details of the identified Variable Methylated Regions (VMRs), the model best explaining their DNAme levels, the variables in said model, and summary statistics. More details about the variables can be found in <https://ericknavarrod.github.io/RAMEN/reference/lmGE.html>.

**Supplementary data 3.** Results of the genome-exposome contribution analysis to methylome variability in PREDO I cord blood samples. The file contains details of the identified Variable Methylated Regions (VMRs), the model best explaining their DNAme levels, the variables in said model, and summary statistics. More details about the variables can be found in <https://ericknavarrod.github.io/RAMEN/reference/lmGE.html>.

**Supplementary data 4.** Results of the genome-exposome contribution analysis to methylome variability in PREDO II cord blood samples. The file contains details of the identified Variable Methylated Regions (VMRs), the model best explaining their DNAme levels, the variables in said model, and summary statistics. More details about the variables can be found in <https://ericknavarrod.github.io/RAMEN/reference/lmGE.html>.

**Supplementary data 5.** List of significant mQTLs found in CHILD cord blood samples (SNP-CpG probe pairs; p-value < 1x10^-5^). Only cis mQTLs (SNP-CpG within Mb) were tested. This file is compressed due to its size.
