## Supplementary figures and tables for "RAMEN: Dissecting individual, additive and interactive gene-environment contributions to DNA methylome variability in cord blood"

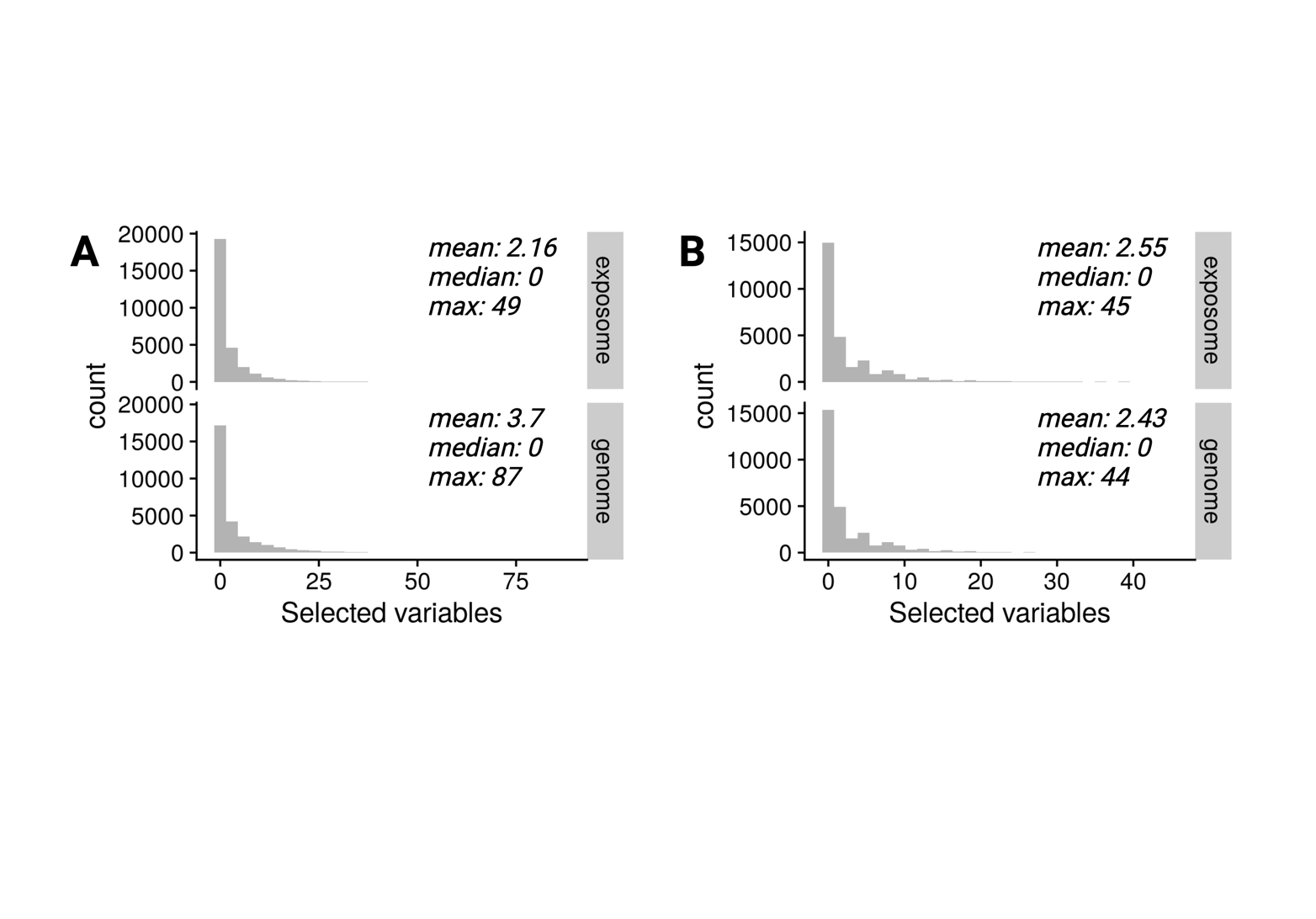

**Figure S1. Distribution of genome and exposome variables selected per VMR under a null association scenario in CHILD.** A) Distribution of selected variables following a permutation on the imbalanced number of genome/exposome variables. B) Distribution of selected variables on a permutated and balanced genome/exposome data set. To balance the data set, following permutation, 94 *cis* SNPs were randomly selected for each VMR (the same number of available exposome variables). VMRs with < 94 *cis* SNPs were excluded from this experiment.

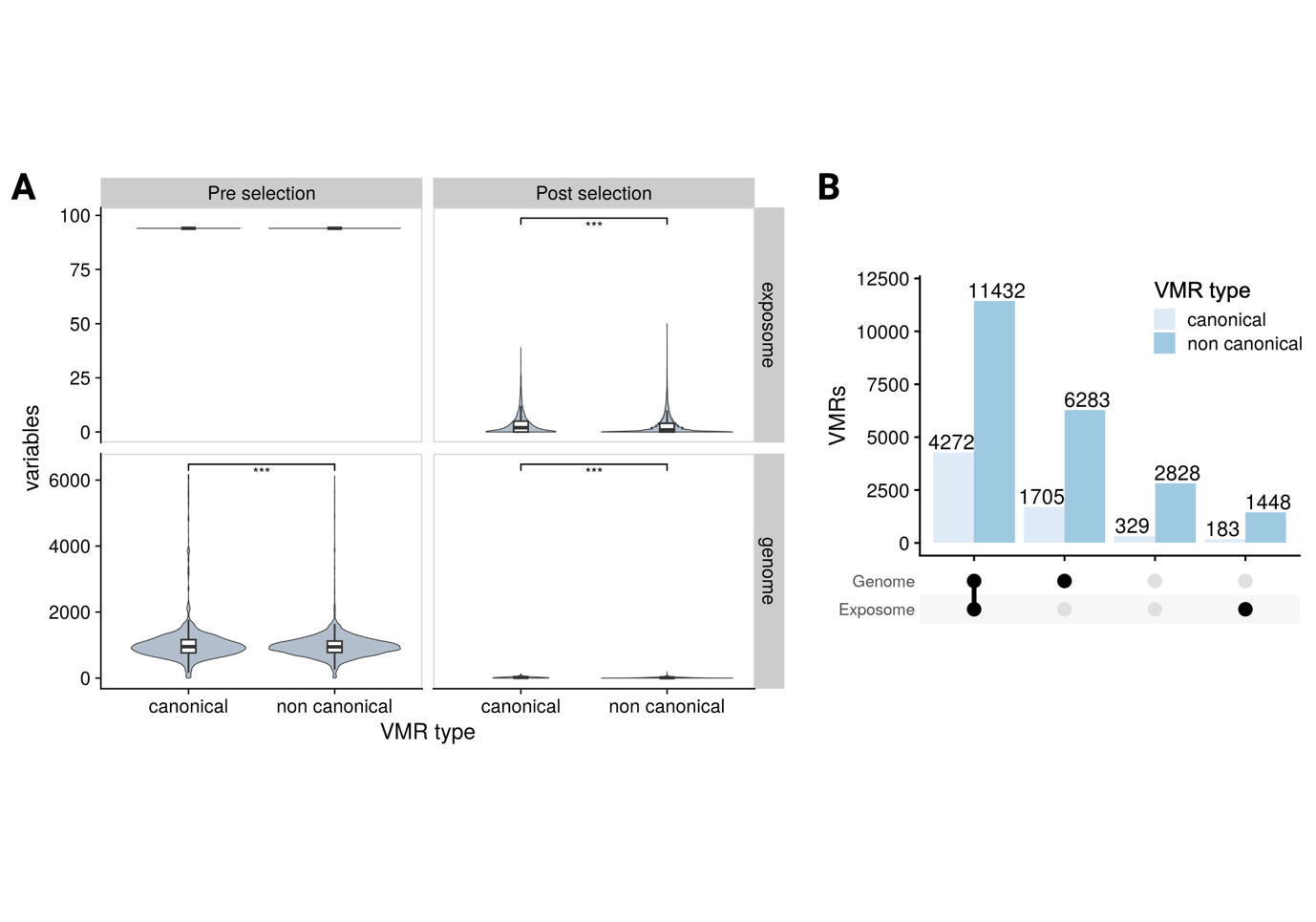

**Figure S2. Differences in variables selection outcomes between canonical and non-canonical VMRs in CHILD.** A) Number of variables before and after RAMEN’s variable selection strategy stratified by canonical and non-canonical VMRs. B) From left to right: number of VMRs with at least one variable selected in both the genome and exposome, only the genome, none, or only the exposome. Bars are stratified by canonical and non-canonical VMRs.

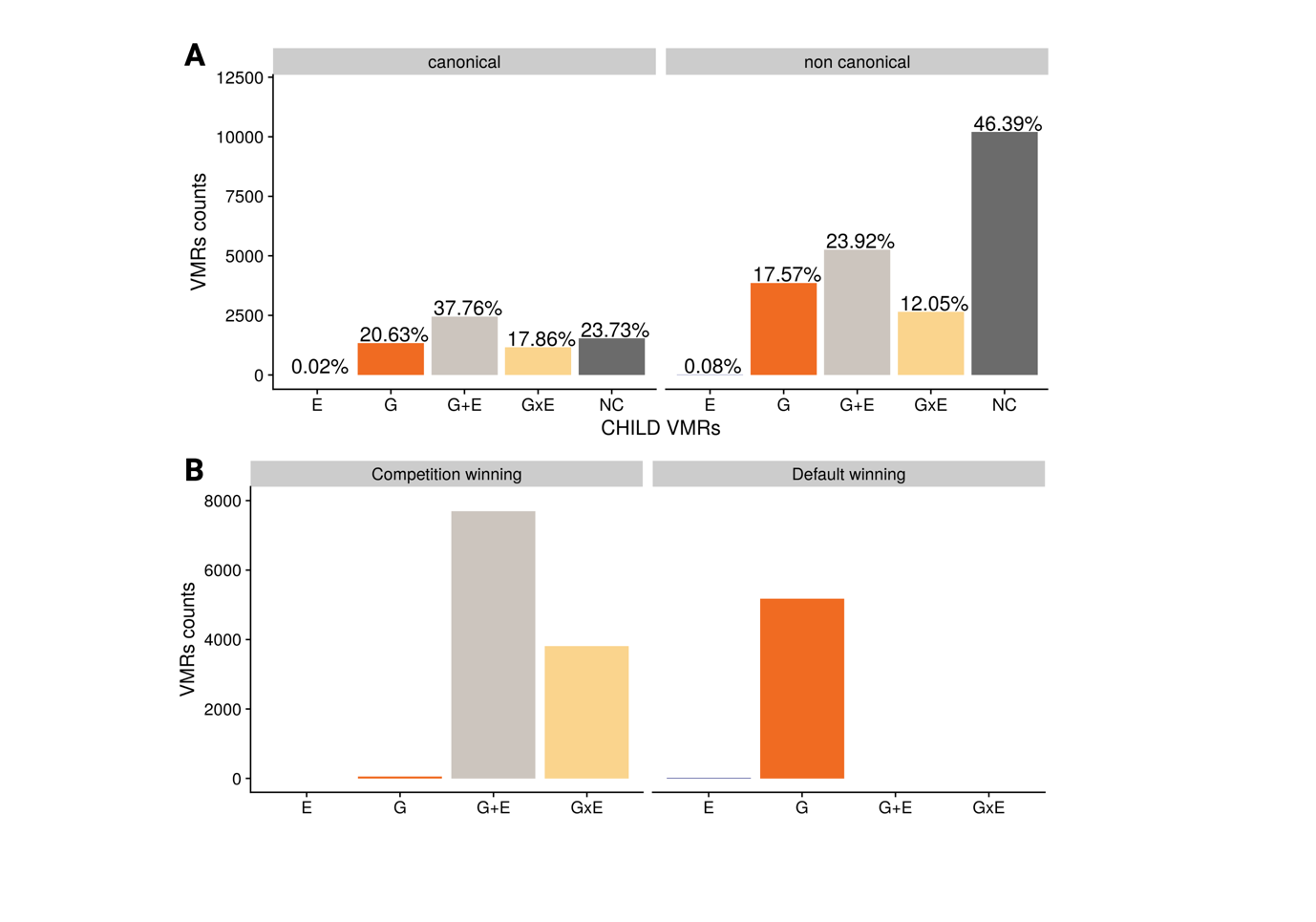

**Figure S3. Number of best VMR models stratified by VMR type and winning model in CHILD.** A) Best models in canonical and non-canonical VMRs. Proportions sum up to 1 within their respective VMR type group. B) Left: VMRs in competition-winning best models, which are models that were selected as winning in the AIC comparison, where they were compared to all possible model categories (G, E, G+E and GxE). These models correspond to the subset of 15,657 VMRs from Figure 3C that passed the permutation analysis threshold. Right: VMRs in default-winning models, which are models that were selected as winning in the AIC comparison only within models of their own group, since there were no variables in the opposite group to fit the full set of the model groups. These models correspond to the subset of 8,035 (only G variables) + 1,630 (only E variables) VMRs from Figure 3C that passed the permutation analysis threshold.

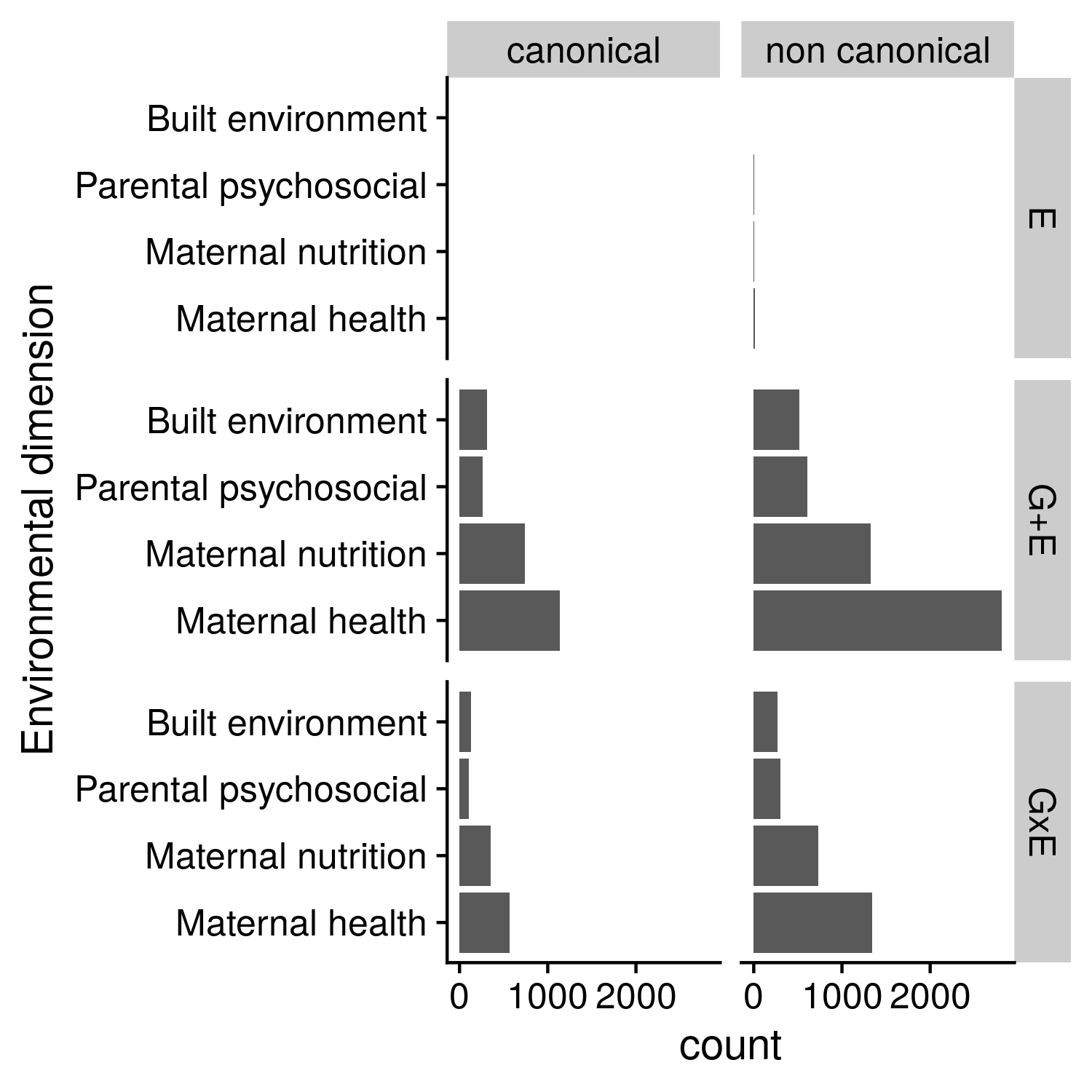

**Figure S4. Prenatal dimension variables in best VMR models in CHILD.**

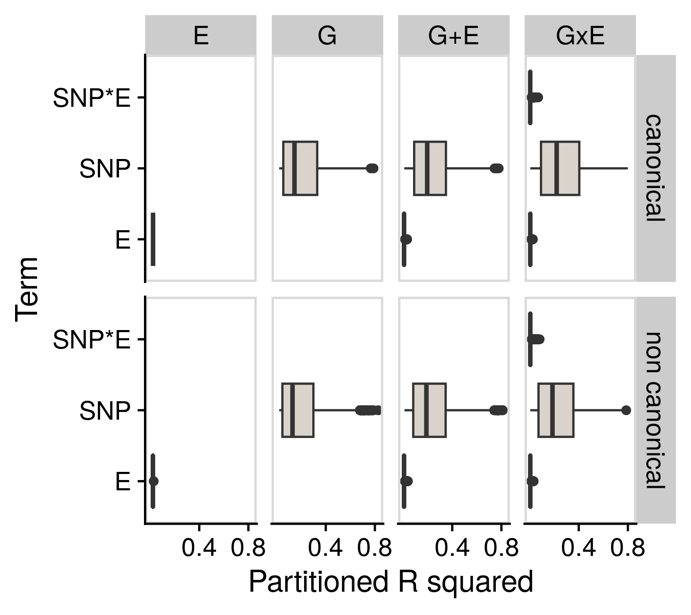

**Figure S5. Decomposed variance in best models in CHILD.** Variance explained by the SNP, SNP*E and E terms across best VMR models, stratified by canonical and non-canonical VMRs.

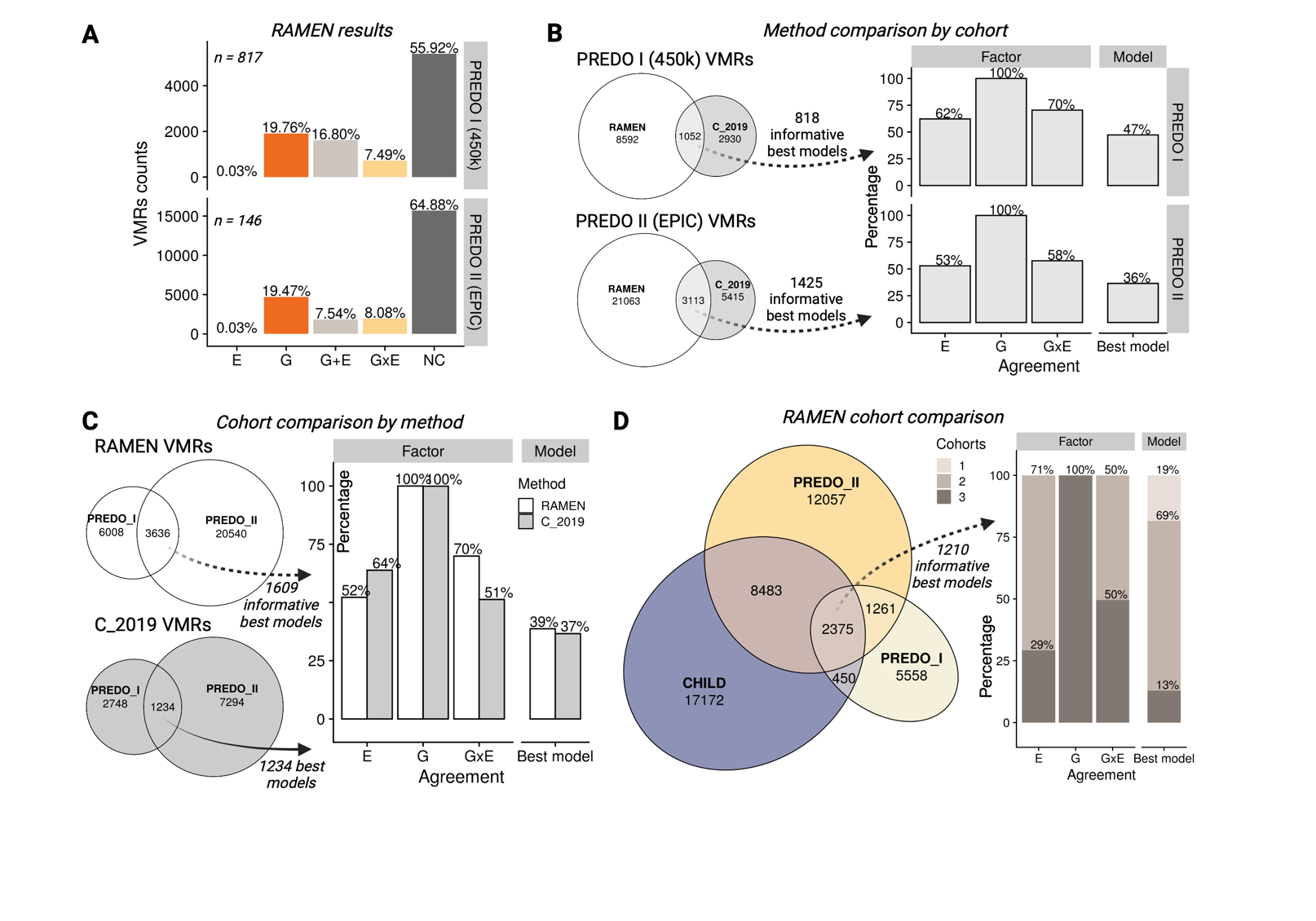

**Figure S6**. Comparison of the model agreements in PREDO I-PREDO II overlapping VMRs using the RAMEN and C_2019 methodologies. The CHILD cohort was not included in this comparison since it was not evaluated in the C_2019 publication. In the C_2019 comparison all models in the overlap are compared because there is not a non-conclusive category.

| Publication | Tissue | DNAme microarray | Cohort | Sample size | Number of VMRs |
| --- | --- | --- | --- | --- | --- |
| Teh *et al.*, 2014 | Umbilical cord | 450k array | GUSTO | 237 | 1,423 |
| Czamara *et al.*, 2019 | Cord blood | 450k array | PREDO I | 817 | 3,982 |
| Czamara *et al.*, 2019 | Cord blood | 450k array | DCHS I | 107 | 6,072 |
| Chatterjee *et al.*, 2021 | Placenta | 450k array | NICHD FGSS | 301 | 5,848 |
| Navarro-Delgado *et al.* | Cord blood | 450k array | PREDO I | 817 | 9,644 |
| Czamara *et al.*, 2019 | Cord blood | EPIC array | PREDO II | 146 | 8,547 |
| Czamara *et al.*, 2019 | Cord blood | EPIC array | DCHS II | 151 | 10,005 |
| Czamara *et al.*, 2019 | Heel prick blood spot | EPIC array | UCI | 121 | 9,525 |
| Navarro-Delgado *et al.* | Cord blood | EPIC array | PREDO II | 146 | 17,578 |
| Navarro-Delgado *et al.* | Cord blood | EPIC array | CHILD | 699 | 28,480 |

**Table S1. Number of Variable Methylated Regions identified in perinatal tissues.** Information extracted from previous studies aiming to estimate the contribution of genetics and environmental factors to DNA methylation variability. The metrics of our study are highlighted in grey.
