## Supplementary methods for "RAMEN: Dissecting individual, additive and interactive gene-environment contributions to DNA methylome variability in cord blood"

**CHILD genotyping pre-processing**

Sample exclusion criteria were the following: call rate £ 0.97, 10% GenCall Score £ 0.4, EPIC-GSA SNP probes mismatches, reported-predicted sex mismatches, detection of sex aneuploidy based on X and Y chromosome signal ratio. SNP exclusion criteria were the following: GenTrain score £ 0.4, cluster separation £ 0.45, call frequency £ 0.97, AB R Mean ≤ 0.4, AB T Mean ≤ 0.2 or ≥ 0.8, AA Frequency = 1 & AA T Mean ≥ 0.2, AA Frequency = 1 & AA T Deviation ≥ 0.04, BB Frequency = 1 & BB T Mean ≤ 0.8, BB Frequency = 1 & BB T Deviation ≥ 0.04, AA Frequency or BB Frequency = 0 & AB T Deviation ≥ 0.5, AB Frequency = 0 & Minor Allele Frequency (MAF) > 0, MAF £ 0.01, Het Excess score £ -0.3 or ³ 0.2 and out of Hardy-Weinberg equilibrium (HWE) (determined through a chi squared test *p* £ 10^-6^).

**CHILD prenatal exposome pre-processing**

Maternal psychosocial dimension

*Psychological questionnaires*

Self-report psychological questionnaires were administered to mothers at the 18^th^ week of pregnancy to measure their retrospective life stress, and concurrent stress, depression symptoms, social support and relationship distress. To score each questionnaire, missing items for individuals that responded ≥70% of the questionnaire were single imputed using predictive mean matching with the *mice* v3.15.0 package in R^1^; individuals that answered the questionnaire but skipped >30% of the items were set as missing in the final score. The questionnaires were processed and scored in the following way:

- Center for Epidemiological Studies – Depression (CES-D): mothers reported the frequency of going through multiple depression behaviours, cognitions and affect during the previous week^2^. Ratings were on a 4-point Likert scale from 0 (none of the time) to 3 (most or all the time). The questionnaire has 20 items where the final CES-D score is obtained by summing each item (with reverse score of items 4, 8, 12 and 16), yielding a score ranging from 0 to 60 with higher scores indicating more depressive symptoms. This is a well-validated measure and with a previously reported Cronbach's α = .87 . In our sample, α = 0.87 and 0.86 for the questionnaires at week 18 and 36 respectively.
- Perceived Stress Scale – 10 items (PSS-10): measures how frequently people find their lives as stressful, unmanageable and overwhelming in the last month^3^. The questionnaire is composed of 10 items that are scored from 0 (never) to 4 (very often). The PSS summary score ranges from 0 to 40 and it is obtained with the sum of every item (with reverse score of items 4, 5, 7 and 8). In our sample, α = 0.9 at week 18 and 36.
- Chronic Stress Response (CSR): measures the amount of stress related to common life conditions and situations. The questionnaire is an adaptation from Wheaton’s 51-item chronic stress inventory^4^, and consists of 37 questions that range from 1 (not true) to 3 (true). Items fall in the dimensions of general stress, work, unemployment, relationships, child care, social life and health. Since the questionnaire has role-specific questions (i.e., some questions only apply if you already have children, are currently in a relationship, employed, etc.), each dimension was scored separately, and the dimension was set as missing if the respondent is not in the corresponding role (there are separate questions in the questionnaire to know the roles). Then, each dimension was standardized and a CSR summary score was obtained by averaging them, ignoring the dimensions that are missing due to not being in that role. The scores cannot be simply added up because the total would be confounded by the number of roles each individual is in. Imputation criteria and procedure here was conducted independently in each dimension. In our sample, α = 0.82.
- Dyadic Adjustment Scale – 10 items (DAS-10): validated standardized questionnaire used to measure the distress in traditional couples^5^. The DAS summary score corresponds to the sum of each item, with items 1,2,5,6,7 and 9 being reverse scored, and it ranges from 0 to 50. Higher scores indicate a more positive dyadic adjustment and a lower level of couple distress. In our sample, standardized α = 0.72.
- Interpersonal Support Evaluation – 12 items (ISEL-12): measures perception of social support in three dimensions: appraisal, belonging and tangible support^6^. The questionnaire is comprised of 12 items that go from 1 (False) to 4 (True). The ISEL-12 summary score consists of an item sum that ranges from 0 to 48, where items 1,2,7,8,11 and 12 are reverse scored. In our sample, α = 0.84.
- Recent Life Events (RLE): negative event checklist based on Turner et al.’s inventory in 1995^7^ that measures the number of stressful life events that happened to the respondent or a close person to them in the last 12 months. Questions include items such as “Was there a serious accident or injury”, “Did a child die”, etc. The checklist consists of 32 events and their start date; the first 23 items ask whether the event has happened to the respondent or to a close person (in separate questions), and the last 9 items ask only about events in the respondent’s life. The answers are binary (0 = no, 1 = yes). For the final score, we summed the number of events that happened to the respondent or to a close person irrespectively on the start date (0 = did not happen, 0.5 = happened to a close person and not to the respondent, 1 = happened to the respondent). The RLE summary score ranges from 0 to 32. In our sample, α = 0.66.

As the CES-D and PSS-10 questionnaires were administered at 18 and 36 weeks of pregnancy, we averaged the summary scores of both periods.

Socioeconomic status

Socioeconomic status metrics aim to capture an individual’s position within a power hierarchy using relatively objective indicators of power, prestige and control over resources^8^. For this study, we used the following SES variables previously processed by the CHILD study team:

- Household income at the 18^th^ week of pregnancy. They were encoded in the following ranges: 1 ($0-$49,999), 2 ($50,000 - $99,999), 3 ($100,000 – $149,999), 4 ($150,000+). Providing ranges helps recall accuracy, since it mitigates participants’ reluctance to disclose “personal information”.
- Maternal education encoded as: 1 (secondary graduation or less), 2 (some post-secondary), 3 (post-secondary graduation), 4 (higher than post-secondary).
- Paternal education encoded as mentioned above.

Maternal and paternal education were maintained as independent variables, since previous studies have found that they can have different effects in children’s outcomes ^9–11^. In addition to the mentioned variables, we created a composed SES variable by taking the first component of a Principal Component Analysis (PCA) using singular value decomposition of the three variables mentioned above.

Built environment dimension

Data linked to the Canadian Urban Environmental Health Research Consortium (CANUE) via subject postal code for the Prenatal 18 week’s visit was retrieved for the following fields: Normalized Difference Vegetation Index (NDVI, which quantifies vegetation greenness), concentration of SO_2_, NO_2_, PM_2.5_, O_3_, temperature, rain and snow.

Maternal nutrition dimension

Dietary intake data were collected using a Food Frequency Questionnaire (FFQ) developed by the Nutrition Assessment Shared Resource (NASR) of Fred Hutchinson Cancer Center, Seattle, Washington^12^. Nutrient calculations were performed by the CHILD study team using the Nutrient Data System for Research (NDSR) software version 2011, developed by the Nutrition Coordinating Center, University of Minnesota, Minneapolis, MN^13^. The processed Daily Intake Dataset fields contain estimated daily intake of more than 130 different nutrients. For this work, only alcohol, caffeine and dietary constituents that mediate 1C metabolism were included, such as folate (B9), other B vitamins (B2, B6, and B12), methionine, choline (betaine) and zinc. Values derived from the FFQ from participants who reported an implausible energy intake in the questionnaire (<500 or >6500 kcal/d) were set as missing. The FFQ was also used to derive the Healthy Eating Index-2010 scores for each component (total fruit, whole fruit, total vegetables, greens and beans, whole grains, dairy, total protein foods, seafood and plant proteins, fatty acids, refined grains, sodium and empty calories), as well as the total HEI score which is a sum of them^14^. Energy adjusted maternal dietary pattern was derived from the FFQ similarly to how it has been previously described in Souza et al. (2016)^15^ using only CHILD data with energy intake values between 500 and 4500 kcal/d^16^. Additional zinc supplementary intake during pregnancy was collected using a separate vitamins and supplement questionnaire at the 18^th^ week of pregnancy.

Maternal health dimension

Maternal health history was collected through questionnaires at week 18 of pregnancy in the following areas: maternal food and pet allergies in the past 12 months, health conditions during pregnancy (e.g., high blood pressure, urinary infection, diabetes, depression, etc.) and smoke exposure during pregnancy. From the allergy questionnaire section, we derived the following variables: allergies reported in > 10 individuals in the open questions (“If other food/pet allergy, specify:”; variables = horse, rabbit, dust and mold), food and non-food related allergies. Additionally, maternal conditions at birth were queried such as bleeding, infections, preeclampsia, hypertension, etc., as well as the birth delivery method. Finally, general maternal characteristics were queried such as gravida, abortions and age.

**Genome-exposome contribution to methylome variability analysis**

LASSO variable selection implementation

Each LASSO model used a tuned penalty parameter λ that minimized the 5-fold cross-validation error (λ_min_) using the *glmnet* R package^17^. We used that value in contrast to the λ within one standard error of the minimum because at this stage the genome and exposome variables were being selected based only on main effects; since we were interested in capturing relevant interactions, we preferred to select a slightly higher number of variables that could potentially have a strong interaction effect when paired with another variable. Furthermore, since we used LASSO as a screening procedure to select variables that were then separately fitted in independent models and compared, the potential issue of overfitting with λ_min_ was not a concern.

Permutation analysis

This procedure explores the explanatory capacity of the Best Explanatory Model just by chance; (i.e. when G and E have no biological association with DNAme). To optimize the number of iterations, we assumed all VMRs to have the same likelihood to exhibit associations under this null scenario. Thus, we were able to pool all the VMR’s R^2^ G/E increment values to estimate the common null distribution and have a high quantity of observations with a small number of iterations. We obtained a bimodal R^2^ G/E increment distribution that we stratified into two groups: marginal (G and E), and joint (G+E and GxE) models. The difference in both groups is explained because the G and E models have a single term, while the rest of the models have 2-3, which increases the R^2^ increment that the later models can have by chance.
